# Keratin 16 spatially inhibits type I interferon responses in stressed and diseased skin

**DOI:** 10.1101/2024.12.27.630544

**Authors:** Erez Cohen, Yang Xu, Sonal Ghodke, Amanda Orosco, Dajia Wang, Craig N. Johnson, Kaylee Steen, Mrinal K. Sarkar, Nurhan Özlü, Lam C Tsoi, Johann E. Gudjonsson, Lucile Marchal, Alain Hovnanian, Carole A. Parent, Pierre A. Coulombe

## Abstract

The stress-induced keratin 16 is broadly used as a biomarker in inflammatory skin disorders while pathogenic variants in *KRT16* cause pachyonychia congenita (PC), a condition in which differentiation and homeostasis are disrupted in palmoplantar epidermis and epithelial appendages. How K16 impacts these disorders at a molecular level is poorly understood. Here we report that K16 spatially restricts type I interferon (IFN) signaling and innate immunity in palmoplantar keratoderma (PPK) lesions in PC patients, imiquimod- and phorbol ester-induced models of sterile inflammation in mouse skin, and poly(I:C)-treated human keratinocytes *ex vivo*. Mechanistically, K16 interacts with effectors of the RIG-I-like receptor (RLR) pathway, including 14-3-3ɛ, and inhibits the 14-3-3ɛ:RIG-I interaction upstream of IFN activation. Topical application of the JAK inhibitor Ruxolitinib reduces the severity of PC-PPK-like lesions in *Krt16 null* mice. These findings uncover a new paradigm for keratin-dependent regulation of innate immunity and suggest a new approach to PC treatment.

**One sentence summary:** *KRT16* negatively regulates type I interferon signaling and innate immune responses in the skin, offering insight into the pathophysiology of inflammatory skin diseases including pachyonychia congenita, psoriasis and others.

## Introduction

Keratinocytes in the interfollicular epidermis constantly encounter environmental, mechanical, and pathological insults. Disruption of keratinocyte homeostasis is linked to many inflammatory diseases, including psoriasis (PSOR), hidradenitis suppurativa (HS), atopic dermatitis, and pachyonychia congenita (PC) (1–5). Over the past decade, research on inflammatory skin diseases has underscored how the interaction between the epidermis and the immune system, known as epithelial-immune crosstalk, is a key determinant for disease onset (6). This communication involves complex interactions between keratinocytes, their niche, and both innate and adaptive immunity (7), and is highly dynamic and tightly regulated. Its failure can disrupt the epithelial barrier (7), while a hyperactive response can trigger inflammatory diseases, such as in the Koebner phenomenon, where keratinocytes stress triggers the resurgence of psoriatic lesions (8). Understanding how keratinocytes manage this crosstalk and recruit innate immune cells, the first immune responders to stress, is critical for the treatment of inflammatory skin diseases.

Keratin intermediate filaments are among the most abundant proteins in skin keratinocytes (9–12) and were recently identified to play a role in the recruitment of neutrophils to stress-responsive interfollicular epidermis (13). Traditionally, keratins have been known to maintain cellular structural integrity in response to various stresses (14, 15). However, recent studies have highlighted their role in regulating cellular signaling including immune response, DNA damage, oxidative stress, and cell growth and differentiation (16–20). It has been suggested that protein-protein interactions are central to keratin function in cellular signaling, with a reliance on post-translational modifications for regulation (21–23). Among the 54 functional keratin genes, three particularly stand out as stress-responsive in skin and other complex epithelia: the type I keratins 16 and 17 (*KRT16*, *KRT17*) and type II keratin 6 paralogs (KRT6A/B/C). These stress-responsive keratins serve as biomarkers for PSOR (24, 25) and many additional chronic skin inflammatory disorders, with expression linked to pathways mediating innate immunity (26). Variants altering the coding sequence of any one of these five keratin genes cause pachyonychia congenita (PC), a debilitating skin disease related to ectodermal dysplasias (2, 27). However, the cellular mechanism and signaling pathways underlying these functions, and the precise significance of these keratins to the onset and development of inflammatory skin diseases, remains underexplored.

Here, we examined the role of K16 in regulating signaling pathways involved in recruitment of innate immunity in both PC and PSOR. K16 is not expressed in healthy epidermis (with the notable exception of palm and sole skin), is rapidly induced in differentiating keratinocytes of epidermis experiencing stresses and is commonly used as a biomarker of PSOR and other chronic inflammatory disorders. *Krt16* null mice, a mouse model for PC, develop spontaneous palmoplantar keratodermas (PPKs), exhibit aberrant differentiation and oxidative stress responses (18, 28) along with increased recruitment of innate immune cells upon chemical irritation (29). Analyses of high-throughput proteomics and transcriptomics datasets uncovered interactions between K16 and members of Type-I Interferon (IFN) pathway, known for its involvement in the onset of PSOR and, specifically, the Koebner phenomenon (30, 31). Although previous reports have identified K16 binding partner, K6, to be upregulated by IFN signaling (32), no known connection between IFN signaling and PC pathophysiology have been investigated to date. This gap is of particular interest as it highlights a potentially targetable pathway involved in PC, a currently untreatable disease.

Informed by our transcriptomic and proteomic analyses, we identified elevated levels of ISG15 and OAS1/OAS3, two type I interferon responsive genes, in the signature palmoplantar keratoderma lesions of individuals with *KRT16* and *KRT6A* mutations. Using *Krt16* null mice, we identified K16 as a negative regulator of imiquimod-induced psoriasiform disease and of the recruitment of innate immune cells to sites of repeated stress. In studies conducted in human N/TERT keratinocytes in culture (33), we show that K16 acts upstream of the IFN pathway activation and limits the interactions between 14-3-3ε and RIG-I, a member of the RLR sensors of dsRNA and LL-37 (34). Application of a FDA-approved, topical JAK inhibitor (JAKi; Ruxolitinib; Opzelura), to PPK lesions of *Krt16* null mice leads to a significant reduction in PPK thickness and IFN effector OAS1 following only 1 week of treatment. Collectively, our findings uncover a new paradigm for the role of keratins in regulating the skin epithelial response to stress and innate immunity, with clear implications for the pathophysiology of inflammatory skin diseases and potential for new therapeutic approaches for PC.

## Results

### K16 expression is limited to the suprabasal epidermis and inversely related to expression of Type I IFN response genes in human disease and mouse models

To explore the potential signaling pathways interacting with K16 in inflammatory diseases, we examined differentially expressed genes in cells with high versus low levels of *KRT16* transcripts using single-cell RNAseq data collected from the skin of PSOR and HS patients (26). These analyses revealed a significant downregulation of IFN signaling genes in *KRT16^high^*expressing cells **(Fig. 1A).** We next examined the spatial relationship between *KRT16* transcripts and those of differentially responsive IFN genes *IFITM3*, *IFITM1*, and *CD74* in spatial single-cell datasets from PSOR and HS skin lesions. These analyses showed that *KRT16* is strongly expressed in the suprabasal layers of epidermis in lesional skin (**Fig. S1A-A’, B-B’)**, consistent with immuno-fluorescence staining for K16 protein **(Fig. 1B-C)** and contrasting with the pattern of K14 and K5 protein **(Fig. S1C-D).** Analysis of spatial single-cell sequencing further shows that the IFN responsive genes *IFITM1*, *IFITM3* and *CD74* are spatially restricted to the basal layer in PSOR and HS (**Fig. S1A-B’’**). To further explore this connection, we screened for K16 interacting proteins by transfecting HaCaT keratinocytes (35) with 3xFLAG-K16 or empty-vector (EV) and performing co-immunoprecipitation, using FLAG antibodies, followed by mass spectrometry **(Fig. 1D)**. In addition to other keratins including K16’s type II keratin partner K6, several IFN response proteins occurred in the filtered list of hits (SAINT>0.8), including RIG-I (DDX58) and MDA5 (IFIH1) members of the RLR (RIG-I-like receptors) family (34) (**Fig. 1D-F)**. To further explore the relationship between *KRT16* and IFN signaling, we examined publicly available microarray (bulk) transcriptomic data from individuals with PC (36). We analyzed data from 6 patients with *KRT16* or *KRT6A-B* mutations (n=3 each; Cao et al., 2015). The effort yielded a significant enrichment for genes involved in type I IFN responses (**Fig. 1G**). Analysis of pathways enriched in the PC-like PPK lesions in the footpad skin of *Krt16* null mice subjected to microarray analysis (28) again showed a differential expression of IFN response genes (**Fig. 1H)**. Together, these observations support the association between K16 and the IFN signaling pathway across inflammatory diseases, model animals and *ex vivo* cell culture models.

**Fig. 1.**
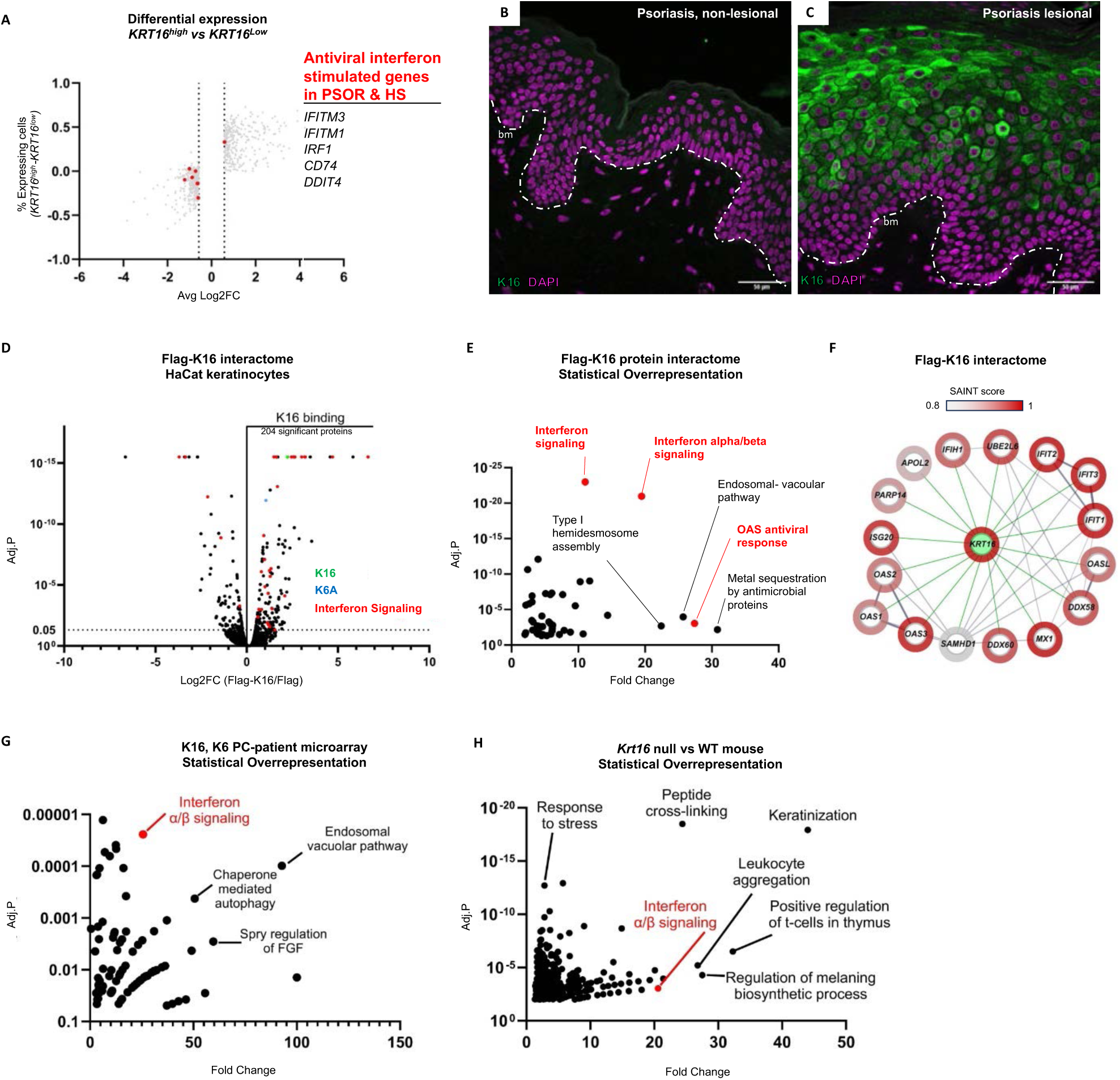
Keratin 16 gene expression and protein interactions are linked to Type I interferon signaling. (A) Analysis of differential expression between *KRT16^high^* and *KRT16^low^* expressing cells in scRNA datasets from PSOR and HS patients. Differentially expressed genes plotted according to “% Expression in the population” (Y axis) vs “Avg Log2” fold change (X axis). Interferon antiviral genes are highlighted in red. (B-C) Immunostaining for K16 (green) and nuclei (DAPI, blue) in tissue sections from non-lesional (B) or lesional skin (C) of individuals with psoriasis. (D) Analysis of IP/MS data from 3xflag-tagged K16 expression in HaCat human keratinocytes. Highlighted are type II keratin K6A (blue) and proteins related to IFN (red). (E) Statistical overrepresentation analysis using Panther (Reactome dataset). Interferon responsive genes and pathways are highlighted in red. (F) Gene network map for highest interacting proteins identified in the IP/MS with SAINT score >0.8. (G) Statistical overrepresentation test for differentially expressed pathways in microarray data from lesional skin of PC patients with mutations in *KRT6* or *KRT16* (source:(36)). (H) Statistical overrepresentation test for differentially expressed pathways in microarray data from lesional footpad skin of *Krt16* null mice (source:(28)).

### Type I interferon responsive genes are elevated in palmoplantar keratoderma lesions of individuals with PC

We obtained biopsies of PPK lesions and non-lesional plantar skin tissue from cases of PC caused by variants in either *KRT16* or *KRT6A* (**Fig. 2A**) to assess the status of type IFN signaling in this disorder. Hematoxylin-eosin staining of sections prepared from these biopsies highlights the severe acanthosis and hyperkeratosis occurring in PC-associated PPK **(Fig. 2B-C’, Fig 2J).** Using indirect immunofluorescence, we first stained for K16 in lesional and non-lesional plantar skin. In non-lesional skin, K16 generally occurred at low levels, with select cells expressing bright K16 immunostaining in the suprabasal layers of the epidermis (**Fig. 2D-D’).** In striking contrast, and in agreement with previous observations in PC patients (29, 36), K16 levels were highly elevated in the suprabasal layers of epidermis in PPK lesions from both PC patients (**Fig. 2E-E’’, K)**. Of interest, we observed aggregates of K16 protein across the suprabasal layers the plantar epidermis in PC biopsies, as previously reported (27, 37–40).

**Fig. 2.**
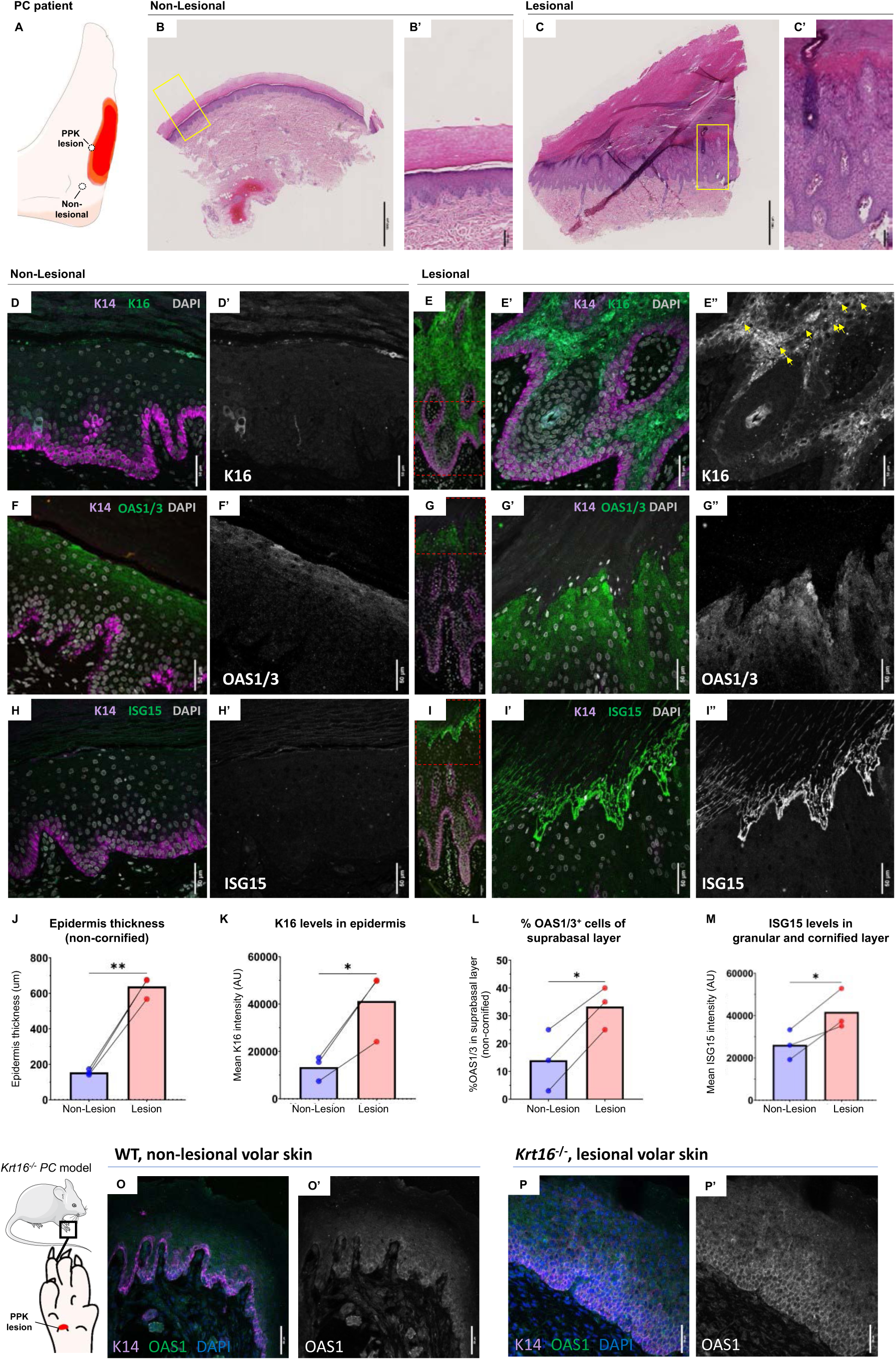
Type I interferon response genes OAS1/3 and ISG15 are elevated in lesional skin of PC patients. (A) Illustration of biopsy sites in plantar skin of PC patients. (B-C’) H&E staining of non-lesional (B) and lesional (C) biopsies from PC patient with K16 pathogenic variant. Yellow box indicate region of inset in (B’,C’). (D-I’’’) Indirect immunostaining for K16 (D-E’’), OAS1/3 (F-G’’) and ISG15 (H-I’’’) in non-lesional (D-D’; F-F’; H-H’) and lesional skin (E-E’’,G-G’’,I-I’’). Channels represent K14 (magenta); nuclei (DAPI, grey); and K16 (D-E’’), OAS1/3(F-G’’) or ISG15 (H-H’’) in green, alongside single-channel images of K16, OAS1/3 or ISG15 respectively. Stitched images PPK lesion full thickness epidermis presented in (E,G,I), red rectangle highlighting the region in (E’,G’,F’). (J-M) Quantification of epidermal thickness (J), K16 levels in suprabasal epidermis (K), % OAS1/3-positive in suprabasal epidermis (L), and ISG15 levels in the upper granular and cornified layer (M) of 3 PC patients with mutations in K16. (O-P’) Indirect immunostaining for OAS1 in volar skin of WT mice (O-O’) and PPK-like lesions of *Krt16* null mice (P-P’). K14 (magenta), OAS1 (green), and nuclei (DAPI, grey) (O,P) presented alongside single-channel image of OAS1 (O’,P’). Scale bars: 1000 μm (A,B), 100 μm (A’,B’), and 50 μm (C-I’’). Fluorescent images represent maximum z-projections across 5 mm depth and quantifications performed on sum projections. Paired student t-test results are reported.

To ascertain the IFN pathway status in both lesional and non-lesional samples, we performed indirect immunostaining for canonical IFN response genes, OAS1/3 and ISG15 (41). Immunostaining for OAS1/3 occurred in the upper granular layers of the epidermis in non-lesional plantar skin **(Fig. 2F-F’)**. This observation is potentially significant, as IFN signaling has been implicated in the development and homeostasis of volar skin (42). Importantly, we observed an increase in the total fraction of epidermis immunopositive for OAS1/3 when comparing non-lesional to lesional skin (**Fig. 2F-G’’;** Patient 1: 33,429um^2^ vs. 168,506um^2^; Patient 2: 44,340um^2^ vs 521,686 um^2^). To account for the increase in epidermal thickness due to acanthosis in the lesional samples, we measured the fraction of OAS1/3-positive area across the entire epidermis (basal to cornified layer). Strikingly, tissue immunopositive for OAS1/3 covers a larger fraction of epidermis even when considering the expansion of suprabasal layer in lesional skin (**Fig. 2F, G, L).** We next assessed whether ISG15 is expressed in non-lesional and PPK lesional plantar skin of individuals with PC. The upper granular layer and cornified layer of non-lesional skin are immunopositive for ISG15 **(Fig. 2H-H’),** extending recent reports of ISG15 function in skin homeostasis (43). The high ISG15 staining intensity near cell-cell borders suggests either ISGylation of membrane-bound, possibly adhesion proteins, or alternatively a secreted pool of ISG15 (43, 44). As seen for OAS1/3, we noticed a significant increase in ISG15 levels in the upper granular layer and in the cornified layers of lesional epidermis from PC patients (**Fig. 2H-I’’, Fig. 2M).** These results further support a role for WT K16 as a regulator of IFN signaling and identify that *KRT16* variants, or dysregulation of K16 protein (as occurs in all forms of PC), lead to elevated IFN response.

To complement these findings, we also performed indirect immunofluorescence for OAS1 in the PPK-like footpad skin lesions in *Krt16^−/−^* mice. While the pattern of staining for OAS1 differs between mouse and human volar skin (compare **Fig. 2O** and **Fig. 2F)**, OAS1 staining is detected in WT footpad epidermis **(Fig. 2O-O’)** and is clearly upregulated in the suprabasal layers of epidermis in lesional, PPK-like footpad skin of *Krt16^−/−^* mice **(Fig. 2P-P’)**. This similarity further supports the notion that K16-dependent regulation of IFN signaling occurs across species and provides a unique opportunity to study the mechanism underlying the association between K16 and IFN across diseases **(Fig. 1)**, and its hyperactivation in PPK lesions of PC patients.

### Loss of *Krt16* leads to exacerbated psoriasiform disease in imiquimod-treated mice

Our transcriptomic and proteomic analysis revealed K16 to associate with IFN signaling across diseases, including PSOR, for which K16 is frequently used as a marker of active disease (24, 25). To directly test whether K16 plays a role in the genesis of psoriasiform lesions, we assessed the response of *Krt16* null mouse skin to imiquimod (IMQ) cream (Aldara™), a widely recognized model of psoriasiform disease (45). *Krt16* null mice and wildtype (WT) litter-mates (FVB strain) were shaved and topically treated for 5 consecutive days with Aldara™ cream, or Vaseline control (**Fig. 3A**), as previously established (45) see Methods). Treated mice were assessed daily **(Fig. 3B-E)**. At baseline, *Krt16* null and WT shaved back skin did not show visually noticeable differences in erythema or scaling, with no difference in thickness or scaling perceptible by touch **(Fig. 3B,D)**. *Krt16* null mice began exhibiting onset of psoriasiform disease as early as day 2 of the 5-day IMQ treatment regimen (**Fig. S2A,B)**. By day 4, a clear exacerbation of disease with increased erythema and scaling was evident relative to controls and WT littermates, **(Fig. 3C,E; Fig. S2G-J).** During tissue harvesting (day 5) we noted an increase in vasculature in IMQ-treated *Krt16* null skin relative to untreated back skin within the same mice (**Fig. S2C-F)**, suggesting that the response is restricted to the treated area. Day 5 skin samples were processed for histological analyses. Both hematoxylin and eosin staining (**Fig. 3F-I)** and indirect immunofluorescence staining (**Fig. 3J-M)** showed increased acanthosis in IMQ-treated *Krt16* null skin compared to IMQ-treated WT controls. As expected, a robust K16 induction was observed in IMQ-treated WT skin but not in controls or *Krt16* null mice **(Fig. 3J-L)**. We also observed an expansion of K14 immunostaining across the suprabasal layers, indicative of aberrant differentiation (26), as previously reported for both the IMQ model in mouse and PSOR patients ((46) and **Fig. S1D)**. We then tested whether loss of K16 leads to changes in cellular proliferation in the IMQ paradigm. We previously showed that the association between induction of K16 and hyperproliferation is a non-cell autonomous phenomenon and takes place at the tissue level (26). Loss of *Krt16* resulted in an increased mitotic index as observed by Ki67 staining after IMQ treatment **(Fig. 3N-Q).** This increase occurs in both the basal and suprabasal layers of the acanthotic epidermis, consistent with the occurrence of aberrant differentiation **(Fig. S2K-K’’)** and previous data from PSOR skin (47).

**Fig. 3.**
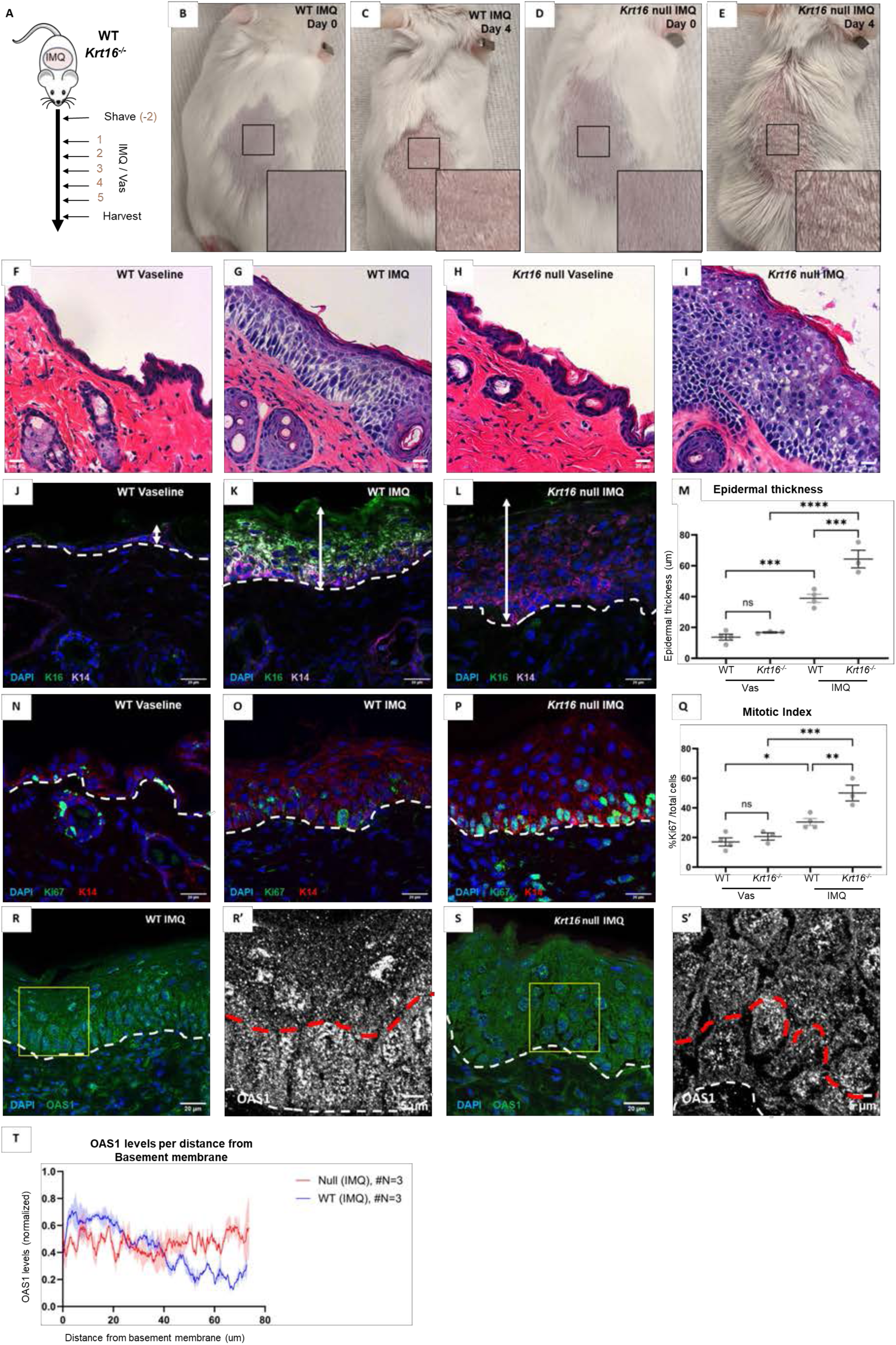
*Krt16* null mice show amplified response to imiquimod model of psoriasiform disease. (A) IMQ application to induce psoriasiform disease in mouse skin (see Methods). (B-E) WT and *Krt16* null mice prior to treatment (day 0; B,D) and following last treatment (day 4; C,E). Insets highlight treated area. (F-I) H&E staining of back skin from WT (F,G) or *Krt16* null mice (H,I) treated with Vaseline control or IMQ. (J-L) Indirect immunofluorescence for K16 (green), K14 (red), and staining for nuclei (DAPI, blue) in back skin sections from WT mice treated with Vaseline (J) or IMQ (K), or *Krt16* null treated with IMQ (L). White dashed delineate basement membrane. (M) Quantification of epidermal thickness across conditions. (N-P) Indirect immunofluorescence for Ki67 (green) and K14 (red), and staining for nuclei (DAPI, blue) in WT mice treated with Vaseline (N), IMQ (O), or *Krt16* null treated with IMQ (P). (Q) Quantification of mitotic index (%Ki67-positive cells) across conditions. (R-S’) Indirect immunofluorescence for OAS1 (green), and staining of nuclei (DAPI, blue) in back skin sections following IMQ treatment of WT (R,R’) and *Krt16* null mice (S-S’). Yellow boxes highlight inset regions (R’,S’) respectively. White dashed lines delineate basement membrane, and red lines depict the interface between basal and suprabasal compartments. (T) Quantification of OAS1 signal intensity (normalized 0-1 per section) from the dermo-epidermal interface (x=0) to the stratum corneum in WT (blue) and *Krt16* null mice (red). Lines represent the mean intensity per um, with standard error across animals shown in light blue or red, respectively. Fluorescent images are maximum projection across 5 mm depth. For (M) and (Q), each dot represents an animal across 3 littermates, One-way ANOVA with Tukey’s multiple comparisons test was used to compare conditions.

Lastly, we tested whether K16 expression negatively correlates with IFN response genes in the IMQ treatment model. To do so we stained for OAS1, which is elevated in PPK lesions of PC patients **(Fig. 2L,P).** Importantly, OAS1 is a robust IFN-responsive gene also known to be elevated in human PSOR lesional skin (**Fig. S1A’’’**; see (48), in psoriasiform disease models (49, 50) and during acute sterile inflammation in mouse skin (13). In line with our analysis of IFN responsive genes in PSOR and HS patients **(Fig. S1A-B’’)**, OAS1 staining was increased in IMQ-treated mice compared to controls (**Fig. 3R-S’, Fig. S2L-M).** In the latter, we noted a clear difference in OAS1 staining in the basal and suprabasal layers of the epidermis, with the basal layer showing higher levels of OAS1 staining **(Fig. 3R-R’,T).** In stark contrast, we did not detect a difference in OAS1 staining between the layers of epidermis in *Krt16* null mice following IMQ treatment **(Fig. 3S-T)**. Of interest, the elevation of OAS1 in the suprabasal layers of *Krt16* null back skin is similar to the findings in the PPK-like lesions spontaneously arising in footpad skin **(Fig. 2O-P)**. These results suggest that K16 participates in restricting the IFN response to the basal layer in stress response and psoriasiform disease.

### Loss of K16 leads to amplified recruitment of neutrophils in a phorbol ester-induced model of acute inflammation

Topical tetradecanoylphorbol-13 acetate (TPA) treatment provides an acute and sterile inflammation model that allows for an assessment of the kinetics and dynamics of neutrophil recruitment to the site of treatment in mouse skin (13), thus complementing the IMQ model. Repeated topical application of TPA results in a <u>T</u>ransiently <u>A</u>mplified neutrophil inflammation <u>R</u>esponse (TAR) in mouse ear skin, which is partially dependent on K17 (13) and is associated with the IFN-responsive genes *OAS1, IFITM1 and IFI16*. We topically applied TPA or Acetone (vehicle control) to mouse ear epidermis of *Krt16* null or WT littermates. We then assessed neutrophil recruitment (Ly6g staining) 6h after a single TPA treatment, or 6h after dual treatment, with the latter showing the characteristic TAR signature **(Fig. 4A-E)**. As we previously showed (13), WT mice showed a modest increase in Ly6g staining 6h after a single TPA treatment, and a markedly accelerated and amplified response at 6h after a second TPA treatment **(Fig. 4B,D)**. By contrast, we observed a significant increase in neutrophil influx in TPA-treated *Krt16* null ear epidermis after both a single or dual TPA treatments **(Fig. 4C,E-G’)**. Otherwise, TPA treatment led to significant thickening of both dermis and epidermis in *Krt16* null relative to WT skin (cartilage to dorsal, **Fig. 4G’).** Surprisingly, the loss of K16 vs. K17 have opposite impacts on the acute response of mouse ear skin to topical TPA.

**Fig. 4.**
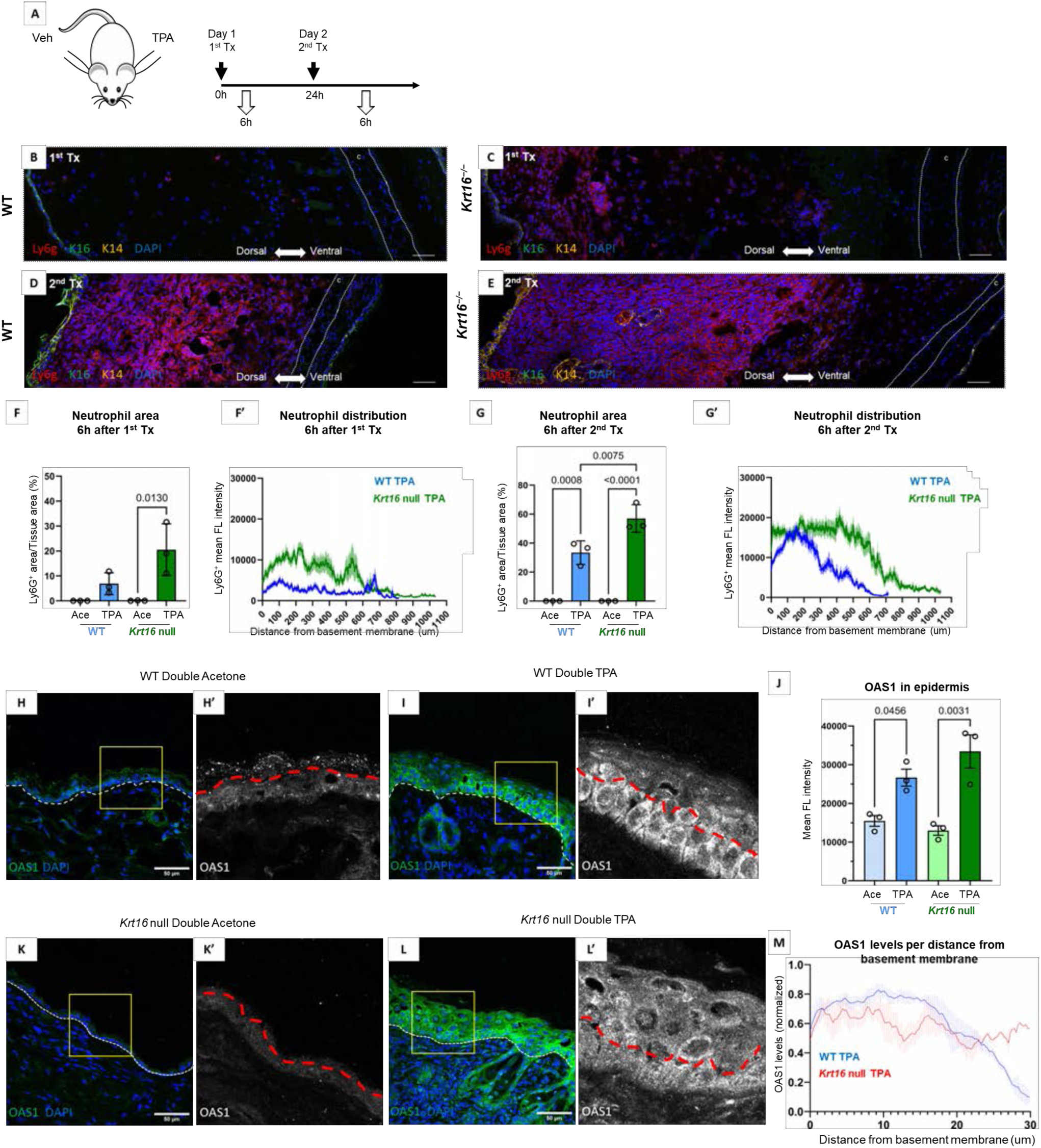
Increased neutrophil recruitment and OAS1 signaling after TPA treatment in Krt16 null mice. (A) Protocol for TPA topical treatment of mouse ear skin (see Methods). (B-D) Indirect immunofluorescence of mouse ear skin sections for neutrophil marker Ly6G (red), K16 (green), K14 (yellow), and staining for nuclei (DAPI, blue), at 6h after single TPA application to WT (B) and *Krt16* null mice (C), or dual TPA application to WT (D) or *Krt16* null ear skin (E). Dashed white lines delineate the cartilage (c) and arrow depicts dorsal to ventral direction. (F-G’) Quantification of neutrophil area (F,G) and distribution (F’,G’) across the dermis of dorsal ear skin at 6h after single TPA treatment (F-F’) or dual TPA treatments (G-G’). (H-I) Indirect immunofluorescence for OAS1 (green), and nuclei staining (DAPI, blue) in ear skin sections after dual acetone (H-H’; K-K’) or dual TPA treatments (I-I’; L-L’). Yellow boxes are magnified in frames (H’, I’, K’, and L’). White dashed line delineates the basement membrane, and red lines depict the interface between basal and suprabasal compartments. (J) Quantification of OAS1 levels in ear epidermis of WT or *Krt16* null mice after dual treatment with Acetone or TPA. (M) Quantification of OAS1 signal intensity (normalized from 0-1 per section) relative to distance from the dermo-epidermal interface (x=0) in WT (blue) and *Krt16* null mice (blue). Lines represent mean intensity per um, with standard error across animals shown in light blue or red respectively. For (F) and (G) each dot represent an animal across 3 littermates with one-way ANOVA with Tukey’s multiple comparisons test used to compare conditions. For (J), each dot represents an animal across 3 littermates, and data from WT and *Krt16 null* TPA used in (M). Holm-Šídák’s multiple comparisons test used to compare conditions in (F).

We next tested whether the loss of *Krt16 in vivo* leads to changes in the basal-restricted IFN response genes normally seen in the TPA-induced TAR paradigm in mouse ear skin. Consistent with previous reports (13), we observed increased OAS1 immunostaining in both WT (**Fig. 4H-J)** and *Krt16* null mouse **(Fig. 4K-L’)** treated twice with TPA, 24h apart. Again, we noted that, in WT mice, the OAS1 induction was significantly higher in the basal relative to the suprabasal layers of treated epidermis **(Fig. 4I’, M).** In line with our observations in PPK lesions and IMQ results, the differential restriction of OAS1 to the basal compartment was clearly lost in *Krt16* null skin **(Fig. 4K-L’)**. These findings further support a role for K16 in inhibiting IFN responses in the suprabasal layer of the epidermis under stress and the downstream recruitment of innate immune cells.

### Loss of *KRT16* in human keratinocyte cells results in amplified responses to poly(I:C), increased recruitment of neutrophils and differentiation defects

To elucidate the molecular mechanism underlying the K16-mediated inhibition of immune recruitment and IFN signaling (**Fig. 5A**), we set out to identify an immortalized human cell line that recapitulates the spatial restriction of K16 expression during keratinocyte differentiation. To do so, we exploited the widely used N/TERT human keratinocyte cell line (33). We adapted the N/TERT 3D culture protocol (51) to generate a post-confluent, uniformly bilayered epithelium in submerged culture under standard KSFM growth medium, without the need for 3D growth at liquid-air interface (see Methods). Under such conditions, expression of keratinocyte differentiation markers, e.g., K10 and filaggrin, is exclusive to the upper layer of keratinocytes (**Fig. S3A-C)**. Further, as is the case in epidermis in situ, we found that whole cell and nuclear size are expanded in the differentiated suprabasal layer of post-confluent N/TERT keratinocytes **(Fig. S3D)**. Importantly, K16 expression is restricted to, and uniform, in the suprabasal layer in this model (**Fig. S3A-B**). We generated and validated a *KRT16* CRISPR KO N/TERT cell line using established methods (52) alongside mock sgRNA controls for the tubulin alpha pseudogene (see Methods and **Fig. 5B-C**).

**Fig. 5.**
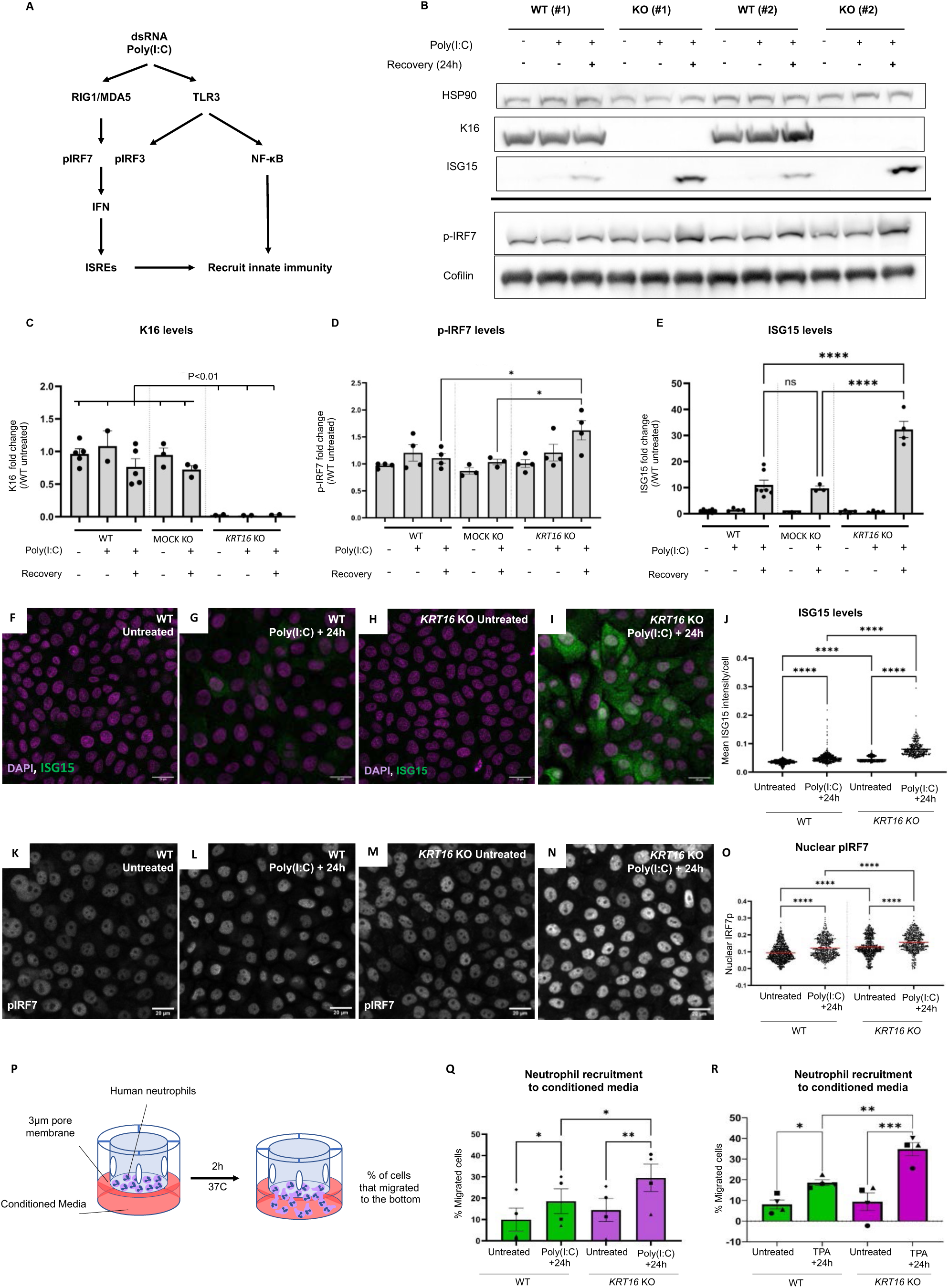
Increased Type I IFN response and neutrophil recruitment in PolyIC-treated *KRT16* null human NTERT keratinocytes. (A) Schematic depicting keratinocyte activation and downstream interferon responses elicited by dsRNA treatment (PolyIC). (B) Western blot of K16, ISG15 and p-IRF7 levels in WT and *KRT16* KO post-confluent N/TERT keratinocyte cultures following (1) untreated, (2) polyIC treatment without recovery and (3) polyIC treatment followed by 24 hours of recovery. Two biological replicates per condition (WT #1, #2; KO #1, #2). HSP90 and Cofilin used as a loading controls. (C-E) Quantification of K16 (C) p-IRF7 (D) and ISG15 (E) levels across polyIC treatment conditions in WT, *KRT16* KO, or MOCK N/TERT keratinocytes. Each dot represents a biological replicate. (F-I) Indirect immunofluorescence for ISG15 (green) and staining for nuclei (DAPI, magenta) in WT (F-G) or *KRT16* KO (H-I) post-confluent N/TERT keratinocyte cultures either untreated (F,H) or at 24h after polyIC treatment (G,I). (J) Quantification of mean ISG15 fluorescence intensity per cell in untreated or polyIC recovered N/TERT keratinocytes. (K-N) Indirect immunofluorescence for p-IRF7 (grey) in WT (K-L) or *KRT16 KO* (M-N) N/TERT keratinocytes untreated (K,M) or at 24 hours after treatment with polyIC (L,N). (O) Quantification of mean nuclear p-IRF fluorescence intensity per cell in untreated or polyIC-recovered N/TERT keratinocytes. (P) Illustration depicting human neutrophil migration assay (see Methods). (Q-R) Quantification of % neutrophil migration towards conditioned media from WT (green) or *KRT16* KO (magenta) in post-confluent N/TERT keratinocytes, untreated or at 24h after treatment with polyIC (Q) or TPA (R). ach dot represents a single biological replicate. One-way ANOVA with Šídák’s multiple comparisons test was used to compare conditions in (C-E). Tukey multiple comparisons test was used to compare conditions in (J) and (O) and data was imaged from 3 technical replicate across 3 biological replicates. For (Q-R), each dot represents a biological replicate of the experiment, utilizing fresh donor neutrophils. Repeated Measures one way ANOVA with Tukey’s multiple comparisons test was used for statistics presented in panels (Q-R).

Utilizing the differentiated, K16-expressing N/TERT model and the *KRT16* KO line, we tested the role of K16 in the induction of IFN signaling through synthetic dsRNA poly(I:C). The latter activates the Type I IFN pathway in keratinocytes (34) and this response is relevant to the skin inflammation observed in PSOR and during the Koebner phenomenon (34). Parental and *KRT16* KO keratinocytes were treated with poly(I:C) for 3h (short response) or 6h with 24h recovery (long response, or “recovery”), and immunostained for upstream regulators of IFN signaling across the RIG-I/MDA5/MAVS pathway, as follows: phosphorylated IRF3 and IRF7 (53); downstream effectors of TLR3 (p65 and phosphorylated p65; see (54); and the canonical IFN response factor ISG15 (55, 56). Poly(I:C) treatment leads to increases in phospho-IRF3 and phospho-p65 within 3h of treatment in both parental and *KRT16 KO* N/TERT cells (**Fig. 5B,D; Fig. S3E-G**). Phosphorylation of these effectors was downregulated following a 24h recovery period, in both genotypes. This is in line with the rapid response expected for both IRF and p65, which is subject to post-translational modifications (57–59). Interestingly, relative to either mock-treated or parental N/TERT cells, we noted a modest yet significant increase in the level of phosphorylated IRF7 in *KRT16* KO cells after a 24h recovery from poly(I:C) treatment **(Fig. 5B,D)**. We next assessed whether ISG15 was amplified in *KRT16* KO cells. At baseline (no treatment), the *KRT16* KO and MOCK KO cell variants show reduced, but not significant, levels of ISG15 relative to parental N/TERT cells (fold change between WT vs KO: 0.64 +/−0.05 SE; WT vs Mock: 0.53 +/−0.25 SE, P>0.1). Such a reduction in IFN signaling has been recently reported for all N/TERT CRISPR KO cell lines at baseline (52). After the 24h recovery from poly(I:C) treatment, however, we noted a strong induction in ISG15 levels **(Fig. 5B,E)**, particularly in the *KRT16 KO* cell line, relative to parental and mock control cells. We further validated these findings using immunofluorescence for ISG15 (**Fig. 5F-J)** and pIRF7 **(Fig. 5K-O).** In line with our *in vivo* observations, therefore, these results establish that loss of *KRT16* leads to an amplification of cellular responses to dsRNA in human keratinocytes *ex vivo*, and that this amplification is cell autonomous.

The amplification of pIRF7 in the *KRT16* KO, together with our data suggesting that effectors downstream of TLR3 (such as phosphorylated P60) are not amplified above WT levels in *KRT16* KO cells, suggested that this response might be dependent on RIG-I/MAVS activation. This was further strengthened by recent reports identifying RIG-I/MDA5 as a key sensor in N/TERT keratinocytes of LL37/dsRNA and downstream activation (60). To address which innate sensor is responsible for induction of the IFN response, we treated WT or *KRT16 KO* N/TERT cell lines with poly(I:C) with inhibitors of RIG-I (RIG012; **Fig. S4A-D**) or TBK1, downstream of MAVS signaling (GSK8612) (**Fig. S4E-H**). We observed that inhibition of RIG-I significantly abrogated ISG15 responses to poly(I:C) in both WT (0.65-fold change, P<0.05) and *KRT16* KO (0.29-fold change) N/TERT cells *ex vivo* as shown in **Fig. S4I**. Notably, *KRT16 KO* cells exhibited greater sensitivity to RIG-I inhibition despite their heightened response to poly(I:C) (53% reduced response in KO cells; p<0.0001; **Fig. S4I)**. Interestingly, TBK1 inhibition led to a significant reduction in the response of *KRT16* KO, but not WT, cells **(Fig. S4J**). This data is in agreement with studies observing TBK1-independent activation of IFN signaling (61), and underscores the reliance of the *KRT16* KO phenotype on the RIG-I/MAVS axis. Finally, to bridge our *ex vivo* and *in vivo* observations, we tested whether the loss of K16 in N/TERT keratinocytes affects their ability to recruit neutrophils after poly(I:C) or TPA treatments. To do so we assessed the ability of conditioned medium collected from treated N/TERT cell cultures to induce the migration of primary human neutrophils utilizing a transwell assay (**Fig. 5P**; see Methods). In response to either poly(I:C) or TPA, loss of K16 led to an amplified recruitment of neutrophils to conditioned media (**Fig. 5Q-R**), further supporting a role for K16 in inhibiting the stress-responsive recruitment of key cellular effectors of innate immunity.

### K16 interacts with 14-3-3ε, a direct regulator of RIG-I receptor activation

To identify the molecular mechanisms underlying the effect of K16 on the response to poly(I:C), we leveraged the endogenous expression of K16 in N/TERT keratinocytes post-confluence to identify K16-interacting proteins using IP/MS in differentiated cells. This screen yielded all known 14-3-3 protein family members (by ranked order: sigma, theta, zeta/delta, epsilon, beta/alpha, gamma and eta; **Supplemental Table 1)**. Follow-up studies were focused on 14-3-3 epsilon(ε) due to its known involvement as an adaptor in RLR signaling and downstream IFN activation (62) and **Fig. 6A**). First, we tested whether loss of K16 impacts the level of 14-3-3ε. No significant change in 14-3-3ε levels could be detected in WT vs. *KRT16* KO cells, whether at baseline or after treatment with poly(I:C) in N/TERT cells **(Fig. 6B).** Using indirect immunofluorescence, we observed colocalization between K16 filaments and 14-3-3ε in the K16-expressing suprabasal layer of post-confluent N/TERT cells (**Fig. 6C-D’’).** To explore this interaction further, we performed proximity ligation assay (PLA) between K16 and 14-3-3ε in poly(I:C)-treated and untreated cells **(Fig. 6E-I)**. Analysis of the PLA signal included both *KRT16* KO cells and the K16-negative basal keratinocytes in parental cells. We measured a significant degree of spatial proximity between K16 and 14-3-3ε in the suprabasal layer of parental N/TERT cells, specifically (**Fig. 6J)**. The number of high-proximity signal was increased further following poly(I:C) treatment **(Fig. 6J)**. To further validate this interaction biochemically, we immunoprecipitated endogenous K16 and analyzed the resulting eluates. Consistent with our IP/MS and PLA data, 14-3-3ε co-immunoprecipitated with K16 in both untreated and poly(I:C) treated N/TERT cells **(Fig. S4K)**.

**Fig. 6.**
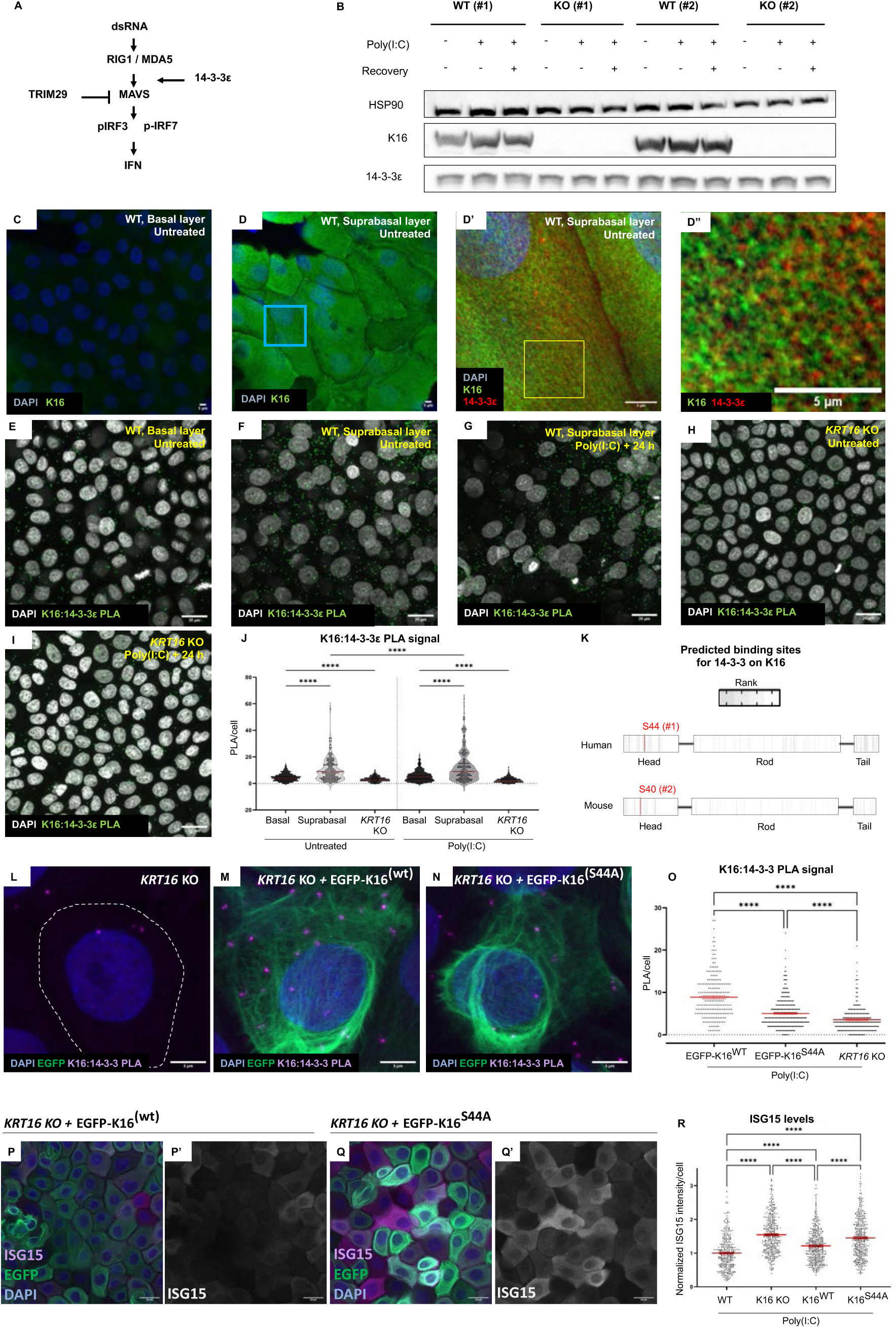
K16 binds 14-3-3epsilon, a regulator of RLR-mediated interferon activation. (A) Schematic depicting the regulation of RIG-I/MDA5 and MAVS signaling by 14-3-3ε and TRIM proteins. (B) Western blot analysis of 14-3-3ε levels in WT and *KRT16* KO post-confluent N/TERT keratinocyte cultures following no treatment (control), polyIC treatment, and at 24h after polyIC treatment (recovery). Two biological replicates per condition are reported (WT #1, #2; KO #1, #2). HSP90 was used for loading control. (C-D) Indirect immunofluorescence for K16 (green) and staining for nuclei (DAPI, blue) in the basal layer (C) and suprabasal layer (D) of post-confluent N/TERT keratinocytes. Blue rectangle in D depict the location of the magnified images in panel D’. (D’-D’’) Airyscan confocal microscopy for K16 (green), 14-3-3ε (red) and nuclei (blue) in suprabasal N/TERT keratinocytes. Yellow indicates areas of colocalization between the two proteins. Yellow rectangle in D’ is magnified in D’’. (E-I) Proximity ligation assay (PLA,green) signal between K16 and 14-3-3ε in basal and suprabasal layers of WT, untreated (E-F) and polyIC treated (F-G), and of *Krt16* KO untreated (H) and polyIC treated (I), N/TERT keratinocytes. Nuclei are stained with DAPI (grey). (J) Quantification of K16:14-3-3ε PLA signal across N/TERT keratinocytes layers, and in *KRT16* KO controls in untreated or at 24h after polyIC treatment. cells were quantified across 3 biological replicates (K) Predicted 14-3-3 binding sites in human and mouse K16. Residues are colored by rank, per the 14-3-3Pred interface (see Methods). Highlighted in red are the S44 and S40 orthologs in human and mouse, and their ranking in parentheses. (L-N) PLA between K16 (Ab #1275) and 14-3-3pan (Ab #H-8) following poly(I:C) treatment in *KRT16 KO* (L), or KO cells stably expressing EGFP-K16^WT^ (M) and EGFP-K16^S44A^ (N) transgenes. PLA depicted in magenta, EGFP in green and nuclei (blue). (O) Quantification of K16:14-3-3 PLA signal across 3 biological replicates. (P-Q’) ISG15 staining following poly(I:C) treatment in *KRT16* KO cells stably expressing EGFP-K16^WT^ (P-P’) and EGFP-K16^S44A^ (Q-Q’) transgenes. ISG15 (magenta), EGFP in green and nuclei (blue) are shown in P,Q. ISG15 is highlighted in grayscale in P’ and Q’. (R) Quantification of ISG15 signal in WT, *KRT16 KO*, and KO cells expressing EGFP-K16^WT^ or EGFP-K16^S44A^ following poly(I:C) treatment.

We next used the 14-3-3pred software (63) to analyze the K16 amino acid sequence for potential 14-3-3 binding sites. 14-3-3 proteins typically bind client proteins in a phosphorylation-dependent manner (64) - this is the case for their previously reported binding to K18 (65), K19 (66), and K17 (20) - a keratin highly homologous to K16. Interestingly, residue S44 in the N-terminal head domain of K16 (orthologous to S40 in mouse), which corresponds to the 14-3-3 binding site in K17 (20), was ranked as the highest predicted site for 14-3-3 interactions **(Fig. 6K)**. To test whether phosphorylation of K16 S44 site is required for 14-3-3 interactions, we generated two stable N/TERT cell lines in the *KRT16* KO background, expressing EGFP fused to either the wildtype form of (EGFP-K16^WT^) or a mutated form of K16 at position 44 (serine to alanine, EGFP-K16^S44A^) **(Fig. S4L)**. We confirmed that these constructs express the full-length fused protein, and that prevailing 14-3-3e levels are not affected by their expression (**Fig. S4L).** We then performed proximity ligation assays across the two lines using K16 and pan-14-3-3 antibody. We observed that S44A mutation reduced the proximity between K16 and 14-3-3 after poly(I:C) treatment compared to EGFP-K16^WT^ (**Fig. 6L-O)**. Consistent with prior studies focused on K17 (67), which shares high homology with K16 head domain, the S44A mutation did not completely abrogate the proximity between K16 and 14-3-3 relative to *KRT16 KO* controls (**Fig. 6L vs Fig. 6N**; **Fig. 6O).** This finding suggests that while phosphorylation at S44 may enhance the affinity between K16 and 14-3-3, it is not required for their interaction. To assess whether K16 S40 phosphorylation occurs *in vivo*, we harvested ear skin from TPA-treated WT mice and used MS to map post-translational modifications across a time span of 0-48h after treatment **(Fig. S4M)**. This effort revealed a continuous increase in K16 protein levels following induction of TPA **(Fig. S4N)** along with increased phosphorylation at the K16 S40 site **(Fig. S4O)**. Together, these results establish that K16 binds 14-3-3ε in human keratinocytes *ex vivo* and establish that the predicted binding site is phosphorylated following TPA application of mouse skin *in vivo*.

To more directly test whether S44 in human K16 plays a role in the amplification of IFN response following poly(I:C) observed in *KRT16* KO, we performed a rescue assay comparing ISG15 levels in WT, *KRT16 KO* cells, and KO cells expressing either the EGFP-K16^WT^ or EGFP-K16^S44A^ mutant, after treatment with poly(I:C). As before, our results identified an amplification in ISG15 levels following poly(I:C) treatment in *KRT16 KO* cells compared to WT controls (54+/−3% increase, P<0.0001, **Fig. 6R).** This amplification was partially rescued by expression of EGFP-K16^WT^ (21+/−4% increase over WT, **Fig. 6P-P’, 6R**). In contrast, expression of EGFP-K16^S44A^ was not sufficient to significantly rescue the amplification of response to poly(I:C) (45+/−3% amplification over WT, n.s. compared to *KRT16* KO alone; **Fig. 6Q-R**). These results support a model where phosphorylation of K16 S44 residue is essential for its role in regulation of IFN in response to dsRNA and, together with our PLA data, supports a role for the 14-3-3: K16 interaction in this paradigm.

### Loss of K16 increases 14-3-3:RIG-I interactions in treated keratinocytes *ex vivo* and in the IMQ model of psoriasiform disease *in vivo*

Given our observation that K16 interacts with 14-3-3ε in N/TERT keratinocytes along with the S44 mutant failure to rescue the *KRT16* null phenotype, we hypothesized that K16 prevents the interaction of 14-3-3ε with RIG-I and the downstream activation of MAVS (62). We first stained N/TERT keratinocytes for RIG-I at baseline and following poly(I:C) treatment **(Fig. 7A-D)**. As expected, our data show increased RIG-I staining in poly(I:C)-treated cells in both parental and *KRT16* KO N/TERT cells. We then assessed spatial proximity between 14-3-3ε and RIG-I using PLA **(Fig. 7E-H)**. We observed an increased 14-3-3ε:RIG-I PLA signal following poly(I:C) treatment in WT keratinocytes (**Fig. 7E-F**), consistent with the known regulation of RIG-I/MAVS via interactions with 14-3-3ε (62). Moreover, we observed an increase in PLA signal in *KRT16* KO keratinocytes relative to parental in both untreated and poly(I:C) treated conditions, suggesting that loss of K16 primes cells towards a heightened response to this treatment **(Fig. 7G-H,M).**

**Fig. 7.**
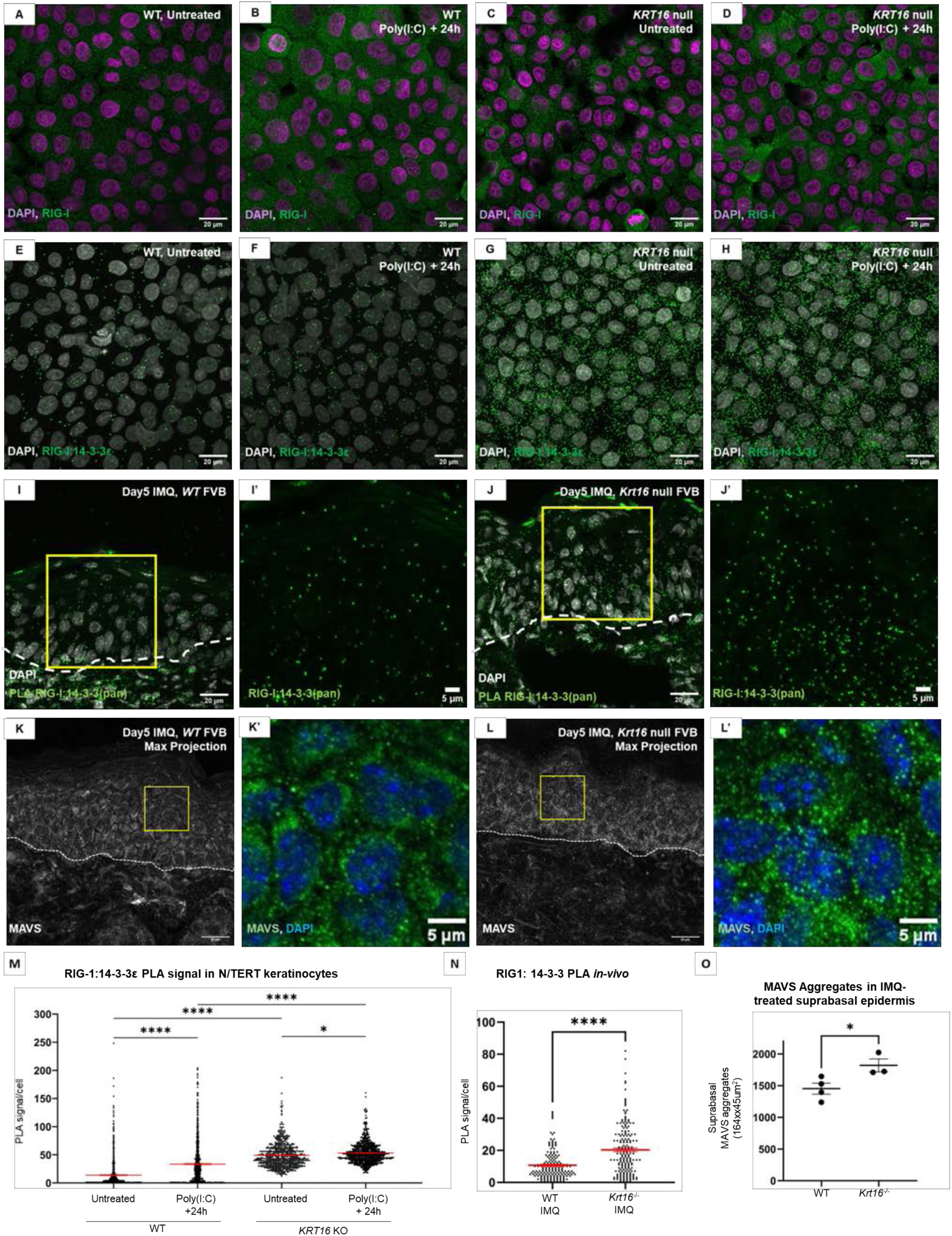
**Loss of K16 leads to increased 14-3-3:RIG-I proximity and MAVS aggregation *ex vivo* and *in vivo.*** (A-D) Indirect immunofluorescence for RIG-I (green) and staining for nuclei (DAPI, magenta) in WT (A-B) and *KRT16* KO (C-D) N/TERT keratinocytes, either untreated (A,C) or at 24h following polyIC treatment (B,D). (E-H) PLA signal probing for close proximity between RIG-I and 14-3-3ε (green) in WT (E-F) or *KRT16* KO (G-H) N/TERT keratinocytes, either untreated (E,G) or at 24h after polyIC treatment (F,H). Nuclei were stained using DAPI (grey). (I-J’) PLA signal probing for close proximity between RIG-I and 14-3-3pan (green) at 5d after IMQ treatment of mouse back skin in either WT (I, inset I’) or *Krt16* null (J, inset J’) mice. Yellow rectangles in I, J depict the location of the magnified image regions in I’, J’. Dashed white lines depict the basement membrane. (K-L’) Indirect immunofluorescence for MAVS at 5 d after IMQ treatment of mouse back skin in WT (K, inset K’) and *Krt16* null (L, inset L’) mice. Yellow rectangles in K, L depict the location of the magnified image regions in K’,L’. Dashed white lines depict the basement membrane. (M) Quantification of RIG:14-3-3ε PLA signal in untreated and at 24h after polyIC treatment in WT or *KRT16* KO N/TERT keratinocytes. (N) Quantification of RIG-I:14-3-3pan PLA signal in the epidermis of IMQ-treated back skin of WT or *Krt16* null mice. (O) Quantification of MAVS aggregates in epidermis of Vaseline- or IMQ-treated back skin of WT or *Krt16* null mice. One way ANOVA with Tukey’s multiple comparisons test was used to compare conditions in (M), and unpaired t-test was used to compare conditions in (N) and (O).

We then tested whether this interaction is impacted by the K16 expression status in the IMQ psoriasiform model *in vivo*. While IMQ is a TLR7/TLR8 agonist and does not directly activate cytosolic RIG-I, recent studies reported on increased levels of RIG-I in keratinocytes after IMQ treatment and identified a mechanistic role for RIG-I in the IMQ-induced production of IL23 by dendritic cells (68). We conducted PLA assays for RIG-I and 14-3-3 in IMQ-treated WT and *Krt16* null mouse skin tissue sections. We found that the PLA signal between RIG-I:14-3-3 is increased in the epidermis of *Krt16* null mice, indicating increased spatial proximity relative to controls (**Fig. 7I-J’,N).** Lastly, we tested whether the loss of K16 has an impact on the aggregation and subsequent activation of MAVS (69) in the IMQ model *in vivo.* Staining for MAVS in skin tissue sections revealed an increased number of aggregates in the epidermis of IMQ-treated WT mice relative to control **(Fig. 7K-L’,O)**. The frequency of MAVS aggregates was significantly increased in *Krt16* null mouse epidermis, however, further supporting a mechanism whereby loss of K16 leads to increased interactions between 14-3-3 and RIG-I and hyperactivation of MAVS and IFN response genes.

### Topical Ruxolitinib treatment attenuates PPK lesion in Krt16 null mouse model of PC

Based on our observations that loss or mutations in K16 lead to amplified IFN responses, we aimed to test whether blocking IFN signaling through inhibition of JAK (JAKi; Ruxolitinib) exerts a therapeutic effect in the *Krt16 null* mouse model of PC. WT or *Krt16* null littermates were topically treated with 35mg control cream (Vanicream; right paw) or topical Ruxolotinib cream (OPZELURA^®^; left paw) twice daily for 7 days as illustrated in **Fig. 8A**. Paws were examined and imaged daily. Treatment with Ruxolitinib or control showed no gross adverse effects either in WT or *Krt16 null* mice **(Fig. 8B-D)**. Gross visual assessment of paws revealed reduced thickness of paw pads with Ruxolotinib treatment, relative to vehicle treatment, in the same individual mice (**Fig. 8B-D, 1 vs 7 days).** Interestingly, we noted that lesion pigmentation was maintained despite the reduced lesion thickness (**Fig. 8D)**. Following 7 days of treatment, mice were euthanized and paws were harvested for histological and molecular assays. Histologically, the thickness of living epidermal layers in PPK lesion was significantly reduced (by ~25%, on average; 411+/−32 μm vs 307+/−36 μm) following Ruxolitinib treatment relative to control cream within the same mice (**Fig. 8E-F’)**. The reduced thickness was not seen under WT (no PPK) conditions, suggesting that the observed reduction in epidermal thickness is specific to the PPK lesion (**Fig. 8J).** Additionally, based on our reports of increased OAS1 expression in the suprabasal layer in PPK lesions from both PC patients (**Fig. 1L)** and *Krt16* null mice **(Fig. 1O-P’)**, we used immunofluorescence to analyze changes in OAS1 across the suprabasal layer of PPK lesions treated with either control or Ruxolitinib cream. First, similar to the histology outcome, we observed no significant change in OAS1 levels in WT mice **(Fig. 8G, K)**. In contrast, PPK lesions in *Krt16* null mice treated with Ruxolitinib showed reduced suprabasal OAS1 levels (1.5 fold reduction, P<0.05, **Fig. 8H-I’, 8K**), indicating that the topical JAKi cream inhibits IFN responsive genes. Together these findings support a role for IFN signaling in the severity of PPK lesions in *Krt16 null* PC mouse model, linking our findings across model systems to a novel potential therapeutic approach for PC.

**Fig. 8.**
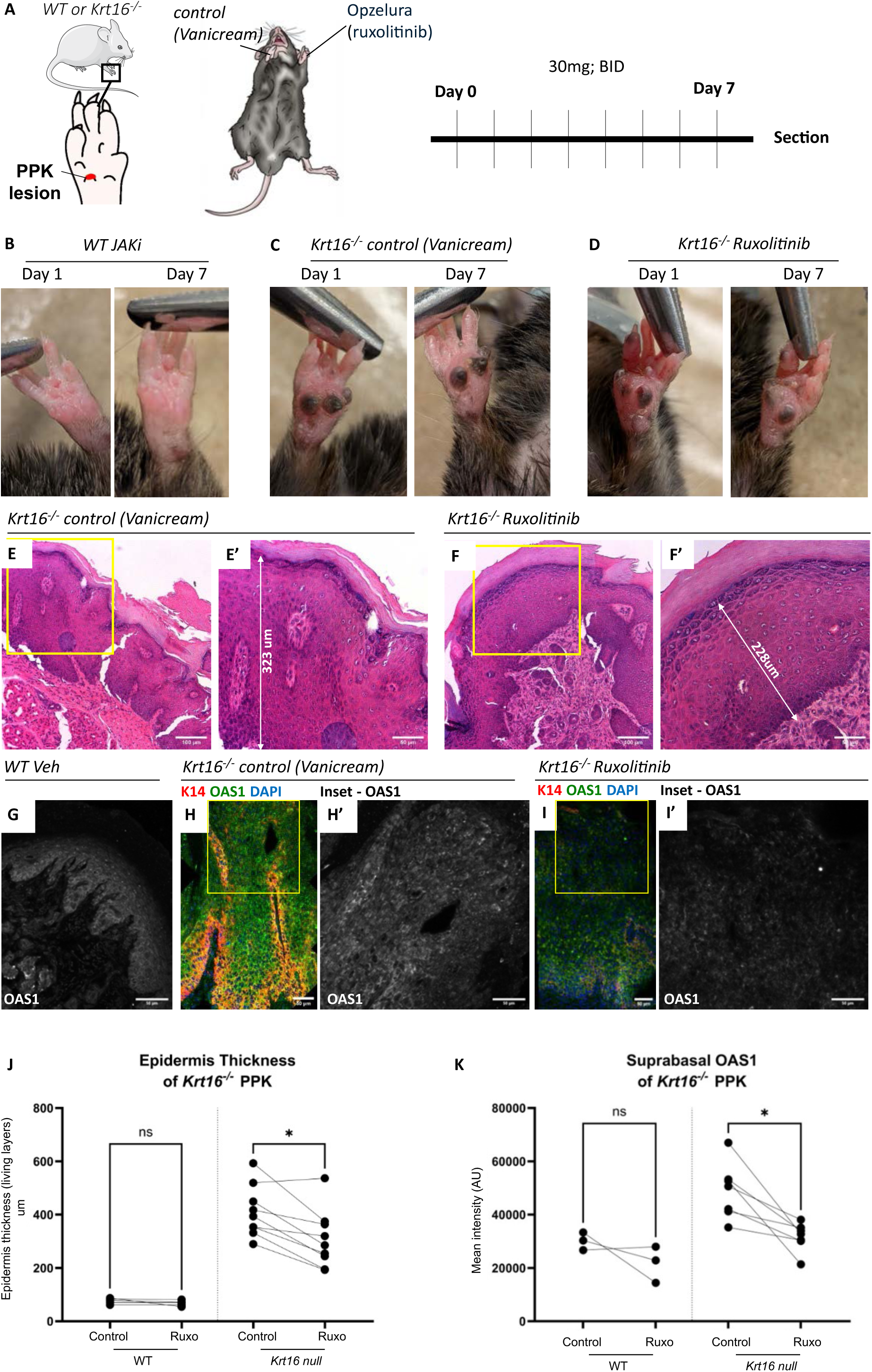
Ruxolitinib treatment of PPK in *Krt16 null* PC mouse model results in decreased lesional thickness and IFN response. (A) Illustration of methodology for applying ruxolitinib or control cream in parallel to *Krt16 null* mice models of PC. (B-D) Images of PPKs from WT (B) or *KRT16 null* mice (C-D) before and after treatment with either control cream (B,C) or ruxolitinib (D). (E-F’) H&E sections from paws of a single individual *Krt16 null* mouse PPKs either treated with control cream (E-E’) or ruxolitinib (F-F’). Yellow border in (E,F) identifies location of respective inset (E’,F’). Arrow identifies living layers of epidermis and their thickness. (G-I’) Indirect immunofluorescence for OAS1 (green), K14 (red) and staining for nuclei (DAPI, blue) in WT (G) and *Krt16* null animals (H-I’) treated with either control cream (H-H’) or at ruxolitinib (I-I’). Full epidermis is presented in panels (G,H,I), with inset location marked with yellow border and corresponding to OAS1 staining in suprabasal layer shown in (H’ and I’). Images (H,I) were stitched from 3 images to showcase the entire epidermis in the PPK lesion. (J-K) quantifications of epidermis thickness (J) or suprabasal OAS1 levels (I) in control or Ruxolitinib (‘Ruxo’) treated WT (no PPK) and *Krt16 null* PPK lesions. Each dot represents a single paw and lines between dots identify the paired paw across individual mice. Experiment was performed across 3 separate litters. Statistical analysis was done using paired one-way ANOVA (REML model) with Tukey’s multiple comparisons test.

## Discussion

The findings reported here identify a role for K16 as a stress-responsive, spatially restricted inhibitor of type I interferon signaling in the suprabasal layers of epidermis. Accordingly, type I IFN signaling is dysregulated in lesional plantar skin of individuals with PC caused by pathogenic variants in *KRT16* or *KRT6A*, and in the PPK-like lesions arising in *Krt16* null footpad skin. In mouse models of both imiquimod-induced psoriasiform disease (back skin) and phorbol ester-induced sterile inflammation (ear skin), loss of *Krt16* results in exacerbated inflammatory phenotypes and loss of IFN spatial restriction—particularly in the suprabasal, differentiating compartment where stress-induced K16 normally occurs. Parallel *ex vivo* studies in differentiated N/TERT human keratinocytes recapitulated mouse model findings and demonstrated that K16 physically interacts with 14-3-3ε, a key upstream regulator of RLR signaling and IFN activation. Our studies establish that K16 physically interacts with 14-3-3ε, a key upstream RLR regulator, in part via residue S44 of K16’s N-terminal head domain. Notably, K16 S44A mutant fails to rescue the amplified response to poly(I:C), supporting a role for phosphorylation at this site in regulating the IFN response. In the Imiquimod-induced psoriasiform mouse model *in vivo,* we show that loss of K16 increases proximity between 14-3-3ε and RIG-I signaling, and aggregation of downstream MAVS proteins, upstream of IFN activation. Motivated by these mechanistic insights, we treated PPK-like lesions in *Krt16* null mice with the FDA-approved topical JAK inhibitor Ruxolitinib, which resulted in significant improvement within one week—underscoring the therapeutic potential of targeting IFN signaling for PPK lesions in PC, an orphan disease currently lacking treatment.

Identifying a role for K16 in regulating interferon signaling ties directly to its use as a biomarker in inflammatory skin diseases showing aberrant IFN signaling, e.g., psoriasis, hidradenitis suppurativa and atopic dermatitis (3, 24, 70). Studies have implicated type I IFN signaling in the keratinocyte response to skin injury (34) and the initiation of skin lesions in vitiligo, psoriasis and lichen planus through the Koebner phenomenon (30, 31). In the Koebner phenomenon, activation of keratinocytes by dsRNA or the antimicrobial peptide LL-37 is thought to trigger the psoriatic autoimmune cascade via upregulation of Type I IFN signaling, specifically through the RIG-I/MDA5 pathway. Interestingly, the regulation of RIG-I/MDA5/MAVS signaling, through 14-3-3ε, was first identified in studies of hepatitis C virus (62). Since then, accumulating evidence has suggested that herpesviruses target 14-3-3ε binding to inhibit IFN signaling (71–73), a mechanism similar to that observed in our study focused on K16. Relevant to the role of K16 across skin development and disease, accumulating evidence suggests significant roles for both IFN and 14-3-3ε in both immunity and skin development diseased skin (74–76). Our findings highlight the tight interplay between K16 expression and RLR and IFN signaling in skin development and disease.

Our observations suggest that high expression of K16 in the context of inflammatory skin diseases may reflect an attempt by keratinocytes to return to a ‘de-activated’ state. In both plantar and interfollicular epidermis, K16 expression is both dynamic and spatially regulated, setting up the stage for its role in immune regulation and keratinocyte differentiation (28, 29, 77). This context-dependent pattern of K16 regulation underscores its crucial role in maintaining epidermal integrity. Consistent with this, mice null for *Krt16* exhibit aberrant differentiation and oxidative stress during development and homeostasis of palmoplantar skin (18, 28, 78). In line with this finding, interferon response, hyperactivation of IRF7, and activation of MAVS have all been linked to oxidative stress and defective Nrf2 responses (79–81). Additionally, recent studies have identified a functional association between dsRNA, RIG-I protein and the expression of K9 (82), a keratin specific to differentiated palmoplantar epidermis that is downregulated upon loss of K16 (28). The prospect of complex interactions between keratinocyte differentiation, oxidative stress, IFN signaling and keratin expression remains to be characterized and could be highly significant for our understanding of palmoplantar skin development and homeostasis, and the formation of PPK lesions in PC and related skin disorders.

Whether K16 regulates similar molecular pathways to promote a ‘de-activated’ state during wound healing, the response to UV, or in additional inflammatory diseases remains to be explored. We previously showed that challenging *Krt16* null mouse skin with injury or topical TPA hyperactivate alarmins (29) including S100A15 (also known as S100A7A or Koebnerisin), which is highly expressed in keratinocytes of psoriatic lesions (83). Of note, neither of the topical treatment paradigms we studied *in vivo* (IMQ, TPA) are direct activators of RIG-I/MDA5/MAVS pathway. Both paradigms, however, are associated with release of damage-associated molecular patterns (DAMPs) (29, 84), activation of IFN responsive genes (13, 50), and regulation by 14-3-3 family of proteins (85). Recent studies have also identified an increase in RIG-I levels in keratinocytes treated with IMQ (68), and showcased LL-37, an activator of RIG-I, as involved in psoriasis (86). While it is possible that abnormal penetration of IMQ or TPA across the skin barrier could drive or exacerbate the phenotype observed in these paradigms, K16 is not expressed in back or ear skin at baseline, and *Krt16* null mice show no detectable abnormalities at these sites before IMQ or TPA treatment. This suggests that increased penetration due to pre-existing barrier defects is unlikely at the onset of treatment. Besides, amplified neutrophil infiltration is already evident in *Krt16* null ear skin after a single TPA treatment. Taken together, these findings indicate that, although altered barrier function resulting from repeated treatments cannot be ruled out, baseline barrier defects are unlikely to account for the initial skin responses observed in our models. To facilitate further investigation into the pathways and mechanisms that may be dysregulated upon loss of K16 across different experimental contexts, we report here on a partial K16 proteome. This dataset identifies several candidate targets and pathways interacting with K16, providing a valuable resource for future studies aimed at elucidating how K16 may alter immune responses in the suprabasal epidermis of stressed skin.

When examining the role of K16 in the context of inflammatory diseases and PC, its relationship with the closely related Type I stress-responsive keratin 17 warrants particular attention. In patients with inflammatory diseases, *KRT16* and *KRT17* are co-expressed in a large population of keratinocytes at the transcriptional level (26). This said, keratinocytes exhibiting a *KRT16^high^*-*KRT17^low^* vs. a KRT16^low^-KRT17^high^ character exhibit distinct transcriptional profiles and are correlated to different immune signaling pathways (26). Similarly, mouse models lacking *Krt16* and *Krt17* display distinct histological features; notably, PPK-like lesions do not form in *Krt17* null mice. Differences can further be observed across patients with PC, where PPKs are significantly more penetrant in PC-K16 subtype than patients with K17 pathogenic variants (IPCC data from 161 patients, 2021). These differences could reflect differential roles and molecular pathways regulated by K16 and K17. In support of the latter, we recently identified a role for K17 in the amplified recruitment of immune responses when the interfollicular epidermis is subjected to repeated stress (13). In this instance, K17 is required for the sustained activation of PKCα in stressed keratinocytes, a mechanism potentially distinct from activation of IFN responses. That K16 and K17, which are highly related in primary structure, would exert opposite influences on immune recruitment in the skin, acting via distinct mechanisms, represents a novel and unanticipated reality.

The differential roles of K16 and K17 might be in part attributed to the distinct biochemical properties of these two keratins, driven by cis-acting determinants located either in the central α-helical rod domain (87, 88), the N-terminal head domain (13) or the C-terminal tail domain (89). Our observation that K16 S40 is phosphorylated after TPA treatment, alongside reduced 14-3-3:K16 proximity and failure of the mutant to rescue the amplified response to poly(I:C) in K16 S44A mutant adds a layer, namely specification via post-translational modifications, to what has been recently described as a “keratin code” (22). How this code is deployed to account for the specific involvement of related keratin proteins that often co-exists in differentiated keratinocytes (e.g., K14, K16, K17) and are connected to distinct signaling pathways and cellular processes now emerges as an issue of fundamental interest. For instance, keratin interactions with 14-3-3 regulate IFN responses (K16 and 14-3-3ε; this study), Hippo signaling (K14, 14-3-3α and YAP1; (19), mTOR signaling (K17 and 14-3-3α; (20, 90)), mitotic progression (K18 and 14-3-3σ; (91) and cell adhesion (K19 and 14-3-3ζ,β,θ; (66)). How the partnership between keratins and 14-3-3 proteins might regulate such a broad variety of pathways and processes, with specificity, provides a significant opportunity to decipher the inner workings of the keratin code. This, in turn, could enable the development of small molecule inhibitors and mimics with promising therapeutic potential.

In the case of PC, therapeutic advances are intimately linked to the ability to decipher the molecular pathways driving its pathology. Previous studies in patients and mouse models have identified several pathways involved in disease progression, including keratinocyte differentiation (28), hyperactivity of EGFR (92), defective Nrf2 signaling and oxidative stress (18). Interestingly, while PC-related PPK lesions have not previously been studied in relation to type I IFN responses, patient bulk RNA-sequencing has identified upregulation of OASL and IGH genes in such lesions (36). Finally, albeit limited in scope (PC-associated PPK is extremely painful and biopsy material is very rare), we here present clear evidence that robust biomarkers of type I IFN responses, e.g., OAS1/3 and ISG15, are significantly elevated in PC lesions. Our findings raise the prospect of testing drugs targeting type I IFN signaling, e.g., JAK inhibitors (93), as a therapeutic strategy of potential interest in PC. Though their use is subject to limitations related to significant side effects (93), such inhibitors have been shown to be effective to treat alopecia areata (94) and related skin disorders (95–99). Unfortunately, even with considerable focus on therapeutic innovation in PC (100), specific, universally effective, or approved treatments remains unavailable. This challenge may, in part, be attributed to the interaction between different pathways involved in PC (100) and their role in lesion development. For example, we have previously reported that use of a small molecule agonist (sulforaphane) to restore Nrf2 function and the cellular redox balance in keratinocytes prevents the formation of PPK-like lesions in the footpad skin of *Krt16* null mice (18). However, unlike our findings with Ruxolitinib, the effectiveness of the Nrf2 activator sulforaphane is reduced once PPK lesions are fully formed (18). A combinatorial approach, aimed at treating various stages of the genesis of PPK lesions (5) using Ruxolitinib, Nrf2 inducers (18, 101) and or the EGF receptor erlotinib (92), might present great therapeutic potential.

Altogether, the findings reported here identify a novel role for K16 in regulating Type I IFN signaling in the context of inflammatory diseases of the skin. Our results provide insights into how keratin expression, and its spatial restriction, can inform epithelial-immune crosstalk. Such insights are highly important for a better understanding of chronic inflammatory skin diseases and exploring new therapeutic approaches for PC.

## Methods

### Study approval and ethics statement

This study utilized histological sections of lesional and non-lesional skin from humans and was approved by the University of Michigan Institutional Review Board (HUM00174864). Participants gave informed consent to participate in a study conducted according to the Declaration of Helsinki Principles. Heparinized whole blood from anonymous healthy human donors was obtained by venipuncture from the Platelet Pharmacology and Physiology Core at the University of Michigan (IRB#HUM00107120). All samples were deidentified and study authors did not have access to the relevant HIPAA information.

Collection of samples from Pachyonychia congenita patients was conducted in accordance with the Declaration of Helsinki and was reviewed and approved by the Internal Review Board (IRB) of the Imagine Institute for Genetic diseases. Patients were seen in the Department of Genomic Medicine for Rare Diseases at The Imagine Institute for Genetic Diseases (Paris, France). All patients and healthy controls were informed and gave their written consent for the use of their remaining biological samples for research purposes. The samples are part of a registered Biobank at INSERM (INSERM DC-2019-3504). Leftover material from skin biopsies was obtained from four patients with pathogenic variants in the keratin 6A (KRT6A) or keratin 16 (KRT16) genes. Biopsy samples originated from calluses of the sole and an adjacent non-lesional area.

### Mouse models and treatment

All experiments were performed on 8-12 weeks old male mice in FVB background and were reviewed and approved by the Institutional Animal Use and Care Committee at the University of Michigan Medical School. *Krt16^−/−^* mouse were previously described and characterized (78, 102). Male mice were exclusively used in the study due to previously reported sexual dimorphism in *Krt16^−/−^* mice (101, 102) relating to temporal fluctuations in the development of PPK in palmoplantar skin. Studies presented in the paper were performed in interfollicular epidermis and observations are thus likely relevant to both sexes. For imiquimod application, *Krt16^−/−^* or WT littermate mice were anesthetized with isoflurane 2 days prior to the experiment (day −2) and had a 60×25mm area of their back skin shaved. Mice were then housed individually for the duration of the experiment. After two days (“day 0”), Imiquimod (5% Aldara™ cream) or Vaseline was applied to the shaved area using a sterile pipette tip under anesthesia. Mice were returned to their housing and treated daily using the same protocol for 4 additional days. At day 5, mice were euthanized and back skin tissue collected for sectioning and analysis. Cryosectioning was performed using CRYOSTAR NX50 Cryostat (Thermo Scientific) and MX35 ultramicrotome blade (Epredia #3053835). Tissues were cut at 5-μm cross-sections and placed on charged microscope slides (VWR #48311–703). Sections were either stained immediately after or stored at −40°C until further use. Application of single or double dose of TPA (Cell Signaling Technology #4174S), or Acetone vehicle control, to dorsal ear has been previously described in detail (13). Briefly, *Krt16^−/−^* or WT littermate mice were treated with single (at 0 h) or double (at 0h and 24h) applications of 20 μl of 0.2 mg/ml of TPA (Cell Signaling Technology), on the dorsal side of ear skin, with the other ear treated with acetone vehicle control. Ear samples were collected at 6h after final application and kept at −40°C for cryosectioning. For treatment using 1.5% ruxolitinib cream (Opzelura™), 12-14 week old C57BL/6 WT or *Krt16 null* littermate mice (male and female) were treated twice a day, with at least 7 hours between treatments, using either Opzelura™ cream or Vanicream® (control). Prior to each treatment, mice were anesthetized for 1 minute and images taken of their palmoplantar keratodermas. 30mg of cream was then applied directly to the paw (Opzelura™ left; Vanicream right) during anesthesia for 1 minute, making sure to rub the cream thoroughly into the plantar skin. The mice were then continued to be anesthetized for an additional 1 minute to allow the cream to soak prior to the mice waking. Apart from a prolonged period of paw licking immediately following the procedure, no adverse effects were observed during the course of treatment. At day 7 of treatment (14 treatments), mice were euthanized, paws were harvested and subjected to either FFPE and histology by the University of Michigan Histology Core, or to cryosectioning and immunofluorescence. A list of mice information is provided as a new **Supplement Table 3.**

### Single cell RNAseq analyses

Single cell RNAseq datasets from human psoriatic lesional skin and HS were originally published by (103–105) and made publicly available at EGAS00001002927 (PSOR), GSE173706 (PSOR) and GSE154775 (HS) GSE154773 (HS). The datasets were reanalyzed using Seurat’s SCTransform “v2” as detailed in (26). Within each dataset, differentially-expressed genes in high vs. low *KRT16*-expressing keratinocytes were identified by comparing top quartiles (“*KRT16*^high^”) vs. lowest quartiles (“*KRT16*^low^”) cells. Differentially expressed genes (adjP<0.05) were then plotted for fold-change and “difference % expression between populations”. Spatial transcriptomics was performed by embedding tissues into a single block, followed by a 5 µm section mounted onto CosMX slide (Nanostring). CosMx spatial library generation and transcriptomics was completed using the manufacturer’s instructions, with tissue sections hybridized with fluorescently labeled oligonucleotide probes targeting specific mRNAs, using the standard custom 1,000 panel.

### Analysis of microarray data from PC patients and *Krt16^−/−^* mouse lesional skin

Microarray datasets were previously published and analyzed (28, 36). For the human PC study (36), gene expression data for patients with K16 and K6 mutations was downloaded, filtered, and analyzed for average p-value across patients. Differentially-expressed genes were analyzed using the Panther (106) and Reactome (107) to identify pathways of significance. For *Krt16^−/−^* mouse microarray dataset, Affymetrix microarray .CEL files from GSE117375 were imported into R using the ‘oligo’ package of Bioconductor.org. Raw Affymetrix probe data were summarized to ENTREZ IDs using version 23 custom Chip Definition Files (CDFs) (108). Data were quantile-normalized using the Robus Multi-Array Average (RMA) technique. The limma package of Bioconductor.org was used to fit array quality weights (109) and linear models with the empirical bayes method to test differences between *Krt16* null and WT samples (110). Significant genes were defined by a Benjamini–Hochberg FDR adjusted P-value of .05 or less and a log2 fold change of 1.5 or greater. Conditional hypergeometric tests were used in the GOstats package or Bioconductor.org, conditioned so that parent terms were not tested with genes of a significant child term.

### Immunoprecipitation and mass spectrometry of cultured keratinocyte proteins

Analysis of the Flag-K16 interactome in cultured HaCat keratinocytes (111) was done in triplicate as follows. HaCat cells were transfected with K16-flag construct or empty vector (EV) control using the nucleofector cell line SE kit, with 500K cells per well and 0.6ug of DNA. Cells were seeded a 6-well plate and allowed to recover for 24h. Cells were then lysed using NETN lysis buffer (150mM NaCl, 1mM EDTA – ethylenediaminetetraacetic acid, 20mM tris buffer pH 8.0 and 0.5% v/v Nonidet P-40) supplemented with Pierce™ Phospho-stop phosphatase inhibitor tablets (Thermofisher Scientific) and cOmplete™protease EDTA-free inhibitor (Millipore Sigma). Samples were bound to the anti-flag M2 magnetic beads (Sigma-Aldrich) and blotted for TCHP, K16 and FLAG to control for pulldown efficiency. Elutes were then submitted to the University of Michigan Proteomics Resource Facility for LC-MS/MS analysis. Data was analyzed with proteome discoverer 2.4 and The Contaminant Repository for Affinity Purification (112) to derive significant interactions (adjP<0.05) and Significance Analysis of INTeractome (SAINT) scores (113). Results were analyzed by String-DB (114) to generate an interactome network between the most highly significant genes (SAINT <0.8). Last, the list of the top 326 significant interacting proteins was subjected to statistical overrepresentation analysis using Panther (106) and Reactome (107) freeware. Significant pathways (FDR adjP<0.05) were retained for consideration.

For immunoprecipitation of endogenous K16 protein from post-confluent, bilayered N/TERT keratinocyte cultures, untreated, Poly(I:C) treated (6h), or Poly(I:C) treated and recovered (6h + 24h recovery) WT and *KRT16* KO cells were lysed using NETN lysis buffer. Samples were assessed for protein concentration using the Pierce BCA assay and 35ug proteins was used as input for the immunoprecipitation. Samples were bound to Dyanbeads protein G (Fisher Scientific) with 2μg anti-K16 antibody (Santa Cruz Biotechnology, #sc-53255), and immunoprecipitated following manufacturer instructions. Elutes were blotted for K16 to control for pulldown efficiency, 14-3-3ε, and cofilin as the input loading control. Last, single replicates of K16 immunoprecipitated elutes (50μg) from untreated and poly(I:C) treated and recovered samples were submitted for University of Michigan Proteomics Resource Facility for LC-MS/MS. Data was analyzed with proteome discoverer 2.4 against SwissProt human protein database and provided as additional supplement (see Supplemental Table 1).

### Mass spectrometry-based analysis of mouse ear skin proteins

Dorsal ear skin samples were collected at 0, 6, 12, 24 and 48h after TPA treatment (4ug, topical). Samples were then resuspended in 8M Urea buffer (50mM Tris-HCl pH 8.5, 8M urea, 1mM EGTA, 1mM PMSF, 2mM DTT) supplemented with Pierce™ Phospho-stop phosphatase inhibitor tablets (Thermofisher Scientific) and cOmplete™ EDTA-free protease inhibitor (Millipore Sigma). After homogenization, lysates were centrifuged at 16,000 × g for 10 min at 4°C and cleared supernatants collected. Samples were boiled with 4× Laemmli sample buffer containing 10% β-mercaptoethanol (BME) at 95°C for 10 minutes and loaded onto an SDS-PAGE gel. Gel pieces from the 50kDa migration area were dissected from the Coomassie-stained gel and digested with Trypsin/Lys-C mix (#PAV5073, Promega) after reduction with 10mM Dithiothreitol and alkylation with 50 mM 2-Chloroacetamine (#C0267, Sigma-Aldrich). The resulting peptides were labeled with TMT10plex Isobaric Label reagents (#90110, ThermoFisher Scientific) according to the manufacturer’s instructions (Labels used:126-0hr; 127N-0hr; 128N-TPA 6h; 129N-TPA 6h; 130N-TPA 12h; 131-TPA 12h; 127C-TPA 24h; 128C-TPA 24h; 129C-TPA 48h; 130C-TPA 48h). Pooled samples were desalted with Sep-Pak solid phase extraction columns. For the phosphorylation analysis, High-Select TiO2 phosphopeptide enrichment kit (A32993, ThermoFisher Scientific) was used according to manufacturer protocol. Protein samples were then processed at the Mass Spectrometry Facility of the Department of Pathology at the University of Michigan, and analyzed using an Orbitrap Ascend Tribrid mass spectrometer (Thermo Fisher Scientific) equipped with an FAIMS source with Vanquish Neo UHPLC. Specifically, MS Samples were analyzed in a 120 min gradient using three compensation voltages (CVs) −40, −55 and −70 V. The full scans (MS1) were acquired over a mass-to-charge (m/z) range of 400-1600 m/z (Orbitrap); the resolution was set to 120K and AGC target 400000; the maximum injection time is 251 ms. Precursor ions with charge states of 2-6 were isolated at 0.7 m/z width and fragmented by HCD (NCE 38%; normalized AGC target of 200%; 45Kresolution; max injection time 50 ms). Dynamic exclusion was 60 seconds. Proteome Discoverer (Thermo Fisher) was used for data analysis and spectra were searched against the Mus musculus UniProt FASTA database (UP000000589). MS1 and MS2 tolerance were set to 10 ppm and 0.6 Da, respectively, and enzyme specificity was set to trypsin with up to two missed cleavages. The search included carbamido-methylation of cysteines and TMT labeling of lysine and N-termini of peptides as fixed modifications; and oxidation of methionine, deamidation of asparagine, glutamine and phosphorylation of serine, threonine, tyrosine as variable modification. Identified proteins and peptides were filtered according to ≤1% FDR threshold.

### Indirect immunofluorescence analysis of mouse and human skin sections

Both mouse and human skin sections were stained using a previously described protocol (Xu et al 2024b). In summary, sections were fixed using 4% paraformaldehyde (PFA, Electron Microscopy Science) for 10 min, washed three times with 1X PBS for 5 minutes, and a circle was drawn around tissue pieces with hydrophobic barrier pen (CALIBIOCHEM #402176). Sections were then blocked with blocking buffer (2% normal goat or donkey serum, 1% bovine serum albumin in 1× PBS) for 30-60 minutes at room temperature. Antibodies diluted in blocking buffer (see appended list of antibodies) were applied in a humid chamber for 2 hours at 37°C. Sections were washed three times with PBS and stained with secondary Alexa-fluor conjugated antibodies (Invitrogen) and DAPI. Last, coverslips (VWR, 1.5 thickness) were mounted using VECTASHIELD® Anti-fade Mounting Medium (Vector Laboratories) for confocal imaging. All sections were imaged for full z-dimensions using a Zeiss LSM800 confocal microscope and analyzed in ImageJ or CellProfiler as detailed in the ‘Image quantification and statistical analyses’ section. Histological analysis using hematoxylin (Thermo Scientific, TA-060-MH) & eosin (Thermo Scientific, 71204) staining was performed per Sigma-Aldrich Procedure No. MHS #1 and images were acquired using a Zeiss fluorescence microscope. For PC patient biopsies, samples were fixed in 4% formaldehyde in PBS (pH 7.4) for 24 hours and then embedded in paraffin. Immunofluorescence and hematoxylin and eosin (HE) staining were performed on 5-µm paraffin sections as described above.

### Post-confluent cultures of human N/TERT keratinocytes and generation of transgenic lines

N/TERT keratinocytes (N/TERT-2G) were grown in Keratinocyte SFM media, supplemented with epidermal growth factor (rEGF, 0.2ng/ml), bovine pituitary extract (BPE, 30 ug/ml), calcium chloride (0.31mM) and penicillin streptomycin (10 units/ml). N/TERT cells were seeded at 300,000 (100mm plate) or at 30,000 cells (6 well plate), with media replaced every 48 h. Cells were monitored daily for confluence and monitored for formation of bilayer. At 4 d post confluence, cells were subjected to treatment (Poly(I:C) or TPA) and processed for imaging, lysis or collection of conditioned media. Generation of CRISPR knock-out (KO) cell lines using non-homologous end joining (NHEJ) via CRISPR/Cas9 and baricitinib was completed by the Functional Analytics CRISPR Core of the University of Michigan Skin Research Center as described (52). Specifically, sgRNA was designed for *KRT16* unique region (5’ GCCTGTCTGTCTCCTCTCGCTTC 3’). In total, 38 clones were generated and tested for *KRT16* DNA and RNA (cDNA Synthesis Kit, Thermofisher Scientific), and proteins levels were assessed by immuno-blotting and staining with K16 antibody. A single clone (#32) showed complete K16 protein KO, secondary to a 4bp deletion and frameshift at amino acid 49, leading to an early stop codon at position 117. To control for non-specific CRISPR phenotypes, CRISPR KO for the tubulin alpha pseudogene was generated referred to as ‘MOCK’ controls, as previously described (115). EGFP-K16^WT^ and EGFP-K16^S44A^ were generated through stable transfection of C3-EGFP-K16 vector (Addgene #6082-1). Vectors were generated through Twist Biosciences and validated using whole plasmid sequencing. Vectors were transfected into 50% confluent N/TERT cultures (0.5ug DNA per vector) using Lonza 4D-Nucleofector X Unit, P3 Primary kit and electroporated utilizing DS-138 setting. Following transfection, cells were grown in G418 media (250 µg/mL) for two weeks and then subjected to FACS sorting using BD FACS Discover S8, to sort individual cells into single colonies based on high EGFP expression. Clones were then grown for additional 2 weeks and imaged for robust and uniform EGFP-keratin staining prior to establishment of stable cell line.

### TPA and PolyIC treatment of N/TERT keratinocyte cultures

For treatment of N/TERT keratinocytes with Polyinosinic-polycytidylic acid sodium salt (poly(I:C), Millipore Sigma), N/TERT keratinocytes were grown to 4 d post-confluence and subjected to: (1) no treatment, (2) 6h treatment with poly(I:C) at 10ug/ml or (3) 6h Poly(I:C) followed by 24h recovery in KSFM media. Cells were fixed with 4% PFA for indirect immunofluorescence or lysed with 6.5M urea buffer (50 mM Tris-HCl pH 7.5, 150 mM NaCL, 5 mM EDTA, 1% Triton X-100, 6.5 M Urea) supplemented with Pierce™ Phospho-stop phosphatase inhibitor tablets (Thermofisher Scientific) and cOmplete™protease EDTA-free inhibitor (Millipore Sigma) for immunoblotting. Both methods were previously described in detail (13). For RIG-I and TBK1 inhibitor studies, cells were first treated for 1 hour with 10uM of inhibitor for RIG-I (RIG012; MCE Cat. No.: HY-147124), TBK1 (GSK8612; MCE Cat. No: HY-111941), or DMSO veh control. Following, poly(I:C) was added for 6 hours, and then replaced with fresh media containing DMSO or inhibitors for 24 hours. Following 24 hours, cells were fixed with 4% PFA and subjected to immunofluorescence protocol as described above. For neutrophil transwell migration assays, conditioned media was collected from untreated or poly(I:C) treated+24h recovered cells using a 3 ml syringe (BD Syringe) with a 0.22 μm PVDF syringe filter (Millipore #SLGVM33RS) and stored at −20°C until use in neutrophil transwell migration assay as previously described (13). For TPA treatment, cells were subjected to (1) 200nM TPA in KSFM media or (2) acetone vehicle control in KSFM media for 6h, followed by a 24h recovery in untreated KSFM media. Conditioned media was then collected from the cells and subjected to neutrophil transwell migration assay.

### Proximity ligation assays

Proximity ligation assays (PLA) were conducted using Duolink PLA Fluorescence kit (Millipore Sigma). In brief, cultured cells or tissue sections were fixed in 4% PFA solution for 20 minutes, and permeabilized with 0.1% Triton X-100 for 10 min at room temperature. Samples were washed 3x with PBS and blocked with either 5% normal goat serum (N/TERT keratinocytes) or 2% normal goat serum, 1% bovine serum albumin (tissue sections) for 16h at 4°C. Primary antibodies were then diluted in blocking buffer and added to sample in a humid chamber for 1h at 37°C. Cells were then washed (1xPBS, 3 times, 5 min) and incubated with PLA probes, ligation mixture and amplification mixture according to the manufacturer’s instructions. Washes with buffers (A, B, 0.01x B) were doubled from manufacturer recommended time and length to improve the signal to noise ratio.

### Image quantification and statistical analyses

Quantitation of immunofluorescence, PLA, and western blots were performed using ImageJ (116) and CellProfiler 4.2.7 (117). Quantification of PLA dots was performed as follows: max z-projection was applied to the images, and primary objects were identified using Otsu two-class threshold for Nuclei (smoothing scale 3; threshold correlation factor 0.5), and Manual threshold strategy for PLA dots (0.05 threshold with 1.344 default smoothing scale). Cells were identified as secondary object based on distance between nuclei (up to 10 μm from the nuclear periphery) using an adaptive Otsu three-class threshold strategy (smoothing scale 10, correlation factor 1). Cells touching the border of the image were discarded from analysis. Outlines of all objects (nuclei, PLA, cells) were saved to provide manual validation and documentation of the analysis. The number of PLA dots per cells was exported and reported. For quantification of immunofluorescence in keratinocytes, sum z-projections were generated for each N/TERT layer and subjected to CellProfiler analysis as described above. Fluorescence intensity for immunostaining (ISG15, pIRF7, RIG-I), and object diameter (nuclei) were calculated per object and results were exported to spreadsheet. Last, to quantify MAVS aggregates, a 165×45um ROI from the suprabasal layer was generated in ImageJ, transformed with max z-projection and used as input for CellProfiler. MAVS aggregates were then identified using a set diameter (5-10 units) and manual thresholding (0.15 threshold with 1.344 default smoothing scale).

For *in vivo* mitotic index analysis, individual cells were counted using ImageJ and categorized based on their Ki67 staining and location with respect to basal or suprabasal layer of epidermis. Mitotic index (Ki67+/total cells) was reported for WT and *Krt16^−/−^*littermates, and analysis of suprabasal mitoses was done in IMQ-treated mice with no suprabasal mitoses observed in Vaseline controls. *In vivo* OAS1 levels were calculated by generating ROI around the epidermis and calculating mean intensity. Additionally, line profiles (width 200pixels) were generated from dermal-epidermal interface to the stratum corneum, and OAS1 signal per distance were recorded. Results across technical replicates (at least 3 lines per section and 3 sections per animal), and animals (at least 3 animals per condition) were aggregated and provided as mean +/− SE in a graphs of OAS1 levels per distance from basement membrane. *In vivo* neutrophil infiltration levels were calculated as the area fraction of Ly6g-positive staining in the TPA-treated skin subtracted by background as previously analyzed (13).

Error bars on histograms represent standard error of the mean (SEM) across biological replicates. All experimental conditions were done with at least 3 biological replicates performed as different batches (separate littermates, separate culture), with at least 2 technical replicates used per biological sample (tissue sections, confocal images, SDS page). Statistical significance was determined by one-way ANOVA followed by Tukey’s multiple comparisons test, Sidak’s multiple comparisons test, and paired or unpaired *t* test. Test information is detailed in the appropriate figure legends. Differences with *p* < 0.05 were considered statistically significant and p values are noted as ns (*p*>0.05), * (*p*<0.05), ** (*p*<0.01), *** (*p*<0.001) and **** (*p*< 0.0001). Graphs generation and all statistical analyses were performed using GraphPad Prism v10.2.2.

### List of antibodies used in study

A list of antibodies and their use is provided in Supplementary Table 2.

## Supporting information

Cohen et al. SUPPL MATERIALS all (Oct 2025)

## Acknowledgments

The authors are grateful to members of the Coulombe and Parent laboratories for advice, to Dr. Venkatesha Basrur and the Proteomics Resource Facility in Department of Pathology at the Univ. of Michigan for mass spectrometry analyses, to the Platelet Pharmacology and Physiology Core at the Univ. of Michigan for providing primary human neutrophils, and to the Skin Biology Center for providing NTERT keratinocytes.

## Author Contributions

EC and PAC together designed the study and analyzed all findings. CAP contributed to the design and interpretation of all experiments involving neutrophils. EC led the studies involving mice and cultured cells with assistance from YX, SG, AO, and DW. KS led the effort to produce a K16 interactome in HaCaT keratinocytes, and NO led analyses involving mass spectrometry. CJ and EC analyzed the single-cell RNAseq datasets, and EC led the effort of integrating them in a shared conceptual framework. MS and JE provided the parental and *KRT16* null N/TERT cell lines, and JE provided key expertise in interferon signaling and chronic inflammatory skin diseases. AH and LM collected biopsies from PC patients, analyzed the histology and provided sections for antigenic analyses. EC and PAC co-wrote the manuscript with input and approval of all co-authors.

## Data and Materials availability

All relevant data is provided within the paper and supporting information files. Single cell RNAseq datasets from human psoriatic lesional skin and HS (Cheng et al., 2018; Gudjonsson et al., 2020; Ma et al., 2023) is publicly available at EGA European Genome-Phenome Archive, EGAS00001002927 (PSOR) or at the Gene Expression Omnibus database GSE173706 (PSOR), GSE154775 (HS), GSE154773 (HS). Spatial transcriptomics data will be made publicly available upon publication of the manuscript.

## References and Notes

1. P. Gallegos-Alcalá, M. Jiménez, D. Cervantes-García, E. Salinas, The keratinocyte as a crucial cell in the predisposition, onset, progression, therapy and study of the atopic dermatitis. International Journal of Molecular Sciences 22, 10661 (2021).

2. S. A. Leachman et al., in *Journal of Investigative Dermatology Symposium Proceedings*. (Elsevier, 2005), vol. 10, pp. 3–17.

3. T. V. Nguyen, G. Damiani, L. A. Orenstein, I. Hamzavi, G. Jemec, Hidradenitis suppurativa: an update on epidemiology, phenotypes, diagnosis, pathogenesis, comorbidities and quality of life. Journal of the European Academy of Dermatology and Venereology 35, 50–61 (2021).

4. A. Rendon, K. Schäkel, Psoriasis pathogenesis and treatment. International journal of molecular sciences 20, 1475 (2019).

5. A. Zieman, P. Coulombe, Pathophysiology of pachyonychia congenita-associated palmoplantar keratoderma: new insights into skin epithelial homeostasis and avenues for treatment. British Journal of Dermatology 182, 564–573 (2020).

6. M. Pasparakis, I. Haase, F. O. Nestle, Mechanisms regulating skin immunity and inflammation. Nature reviews immunology 14, 289–301 (2014).

7. D. Rosenblum, S. Naik, Epithelial–immune crosstalk in health and disease. Current opinion in genetics & development 74, 101910 (2022).

8. G. Weiss, A. Shemer, H. Trau, The Koebner phenomenon: review of the literature. Journal of the European Academy of Dermatology and Venereology 16, 241–248 (2002).

9. R. Moll, M. Divo, L. Langbein, The human keratins: biology and pathology. Histochemistry and cell biology 129, 705–733 (2008).

10. E. Cohen, C. Johnson, C. J. Redmond, R. R. Nair, P. A. Coulombe, Revisiting the significance of keratin expression in complex epithelia. Journal of Cell Science 135, jcs260594 (2022).

11. T. Sun, H. Green, Keratin filaments of cultured human epidermal cells. Formation of intermolecular disulfide bonds during terminal differentiation. Journal of Biological Chemistry 253, 2053–2060 (1978).

12. X. Feng, H. Zhang, J. B. Margolick, P. A. Coulombe, Keratin intracellular concentration revisited: implications for keratin function in surface epithelia. The Journal of investigative dermatology 133, 850 (2012).

13. Y. Xu, E. Cohen, C. N. Johnson, C. A. Parent, P. A. Coulombe, Repeated stress to the skin amplifies neutrophil infiltration in a keratin 17-and PKCα-dependent manner. PLoS biology 22, e3002779 (2024).

14. E. Fuchs, D. W. Cleveland, A structural scaffolding of intermediate filaments in health and disease. Science 279, 514–519 (1998).

15. M. B. Omary, P. A. Coulombe, W. I. McLean, Intermediate filament proteins and their associated diseases. New England Journal of Medicine 351, 2087–2100 (2004).

16. R. P. Hobbs, J. C. Lessard, P. A. Coulombe, Keratin intermediate filament proteins–novel regulators of inflammation and immunity in skin. Journal of cell science 125, 5257–5258 (2012).

17. R. R. Nair et al., A role for keratin 17 during DNA damage response and tumor initiation. Proceedings of the National Academy of Sciences 118, e2020150118 (2021).

18. M. L. Kerns et al., Oxidative stress and dysfunctional NRF2 underlie pachyonychia congenita phenotypes. The Journal of clinical investigation 126, 2356–2366 (2016).

19. Y. Guo et al., Keratin 14-dependent disulfides regulate epidermal homeostasis and barrier function via 14-3-3σ and YAP1. Elife 9, e53165 (2020).

20. S. Kim, P. Wong, P. A. Coulombe, A keratin cytoskeletal protein regulates protein synthesis and epithelial cell growth. Nature 441, 362–365 (2006).

21. P. Li, K. Rietscher, H. Jopp, T. M. Magin, M. B. Omary, Posttranslational modifications of keratins and their associated proteins as therapeutic targets in keratin diseases. Current Opinion in Cell Biology 85, 102264 (2023).

22. J. Di Russo, T. M. Magin, R. E. Leube, A keratin code defines the textile nature of epithelial tissue architecture. Current Opinion in Cell Biology 85, 102236 (2023).

23. H.-M. Pallari, J. E. Eriksson, Intermediate filaments as signaling platforms. Science’s STKE 2006, pe53–pe53 (2006).

24. S. Jiang, T. E. Hinchliffe, T. Wu, Biomarkers of an autoimmune skin disease—psoriasis. Genomics, Proteomics and Bioinformatics 13, 224–233 (2015).

25. R. Rashmi, K. Rao, K. Basavaraj, A comprehensive review of biomarkers in psoriasis. Clinical and experimental dermatology 34, 658–663 (2009).

26. E. Cohen et al., Significance of stress keratin expression in normal and diseased epithelia. Iscience 27, (2024).

27. W. McLean et al., Keratin 16 and keratin 17 mutations cause pachyonychia congenita. Nature genetics 9, 273–278 (1995).

28. A. G. Zieman, B. G. Poll, J. Ma, P. A. Coulombe, Altered keratinocyte differentiation is an early driver of keratin mutation-based palmoplantar keratoderma. Human molecular genetics 28, 2255–2270 (2019).

29. J. C. Lessard et al., Keratin 16 regulates innate immunity in response to epidermal barrier breach. Proceedings of the National Academy of Sciences 110, 19537–19542 (2013).

30. X. Zhang et al., Characteristics and pathogenesis of Koebner phenomenon. Experimental dermatology 32, 310–323 (2023).

31. L.-j. Zhang, Type1 interferons potential initiating factors linking skin wounds with psoriasis pathogenesis. Frontiers in immunology 10, 1440 (2019).

32. Y. Zhao, U. Gartner, F. J. Smith, W. I. McLean, Statins downregulate K6a promoter activity: a possible therapeutic avenue for pachyonychia congenita. Journal of Investigative Dermatology 131, 1045–1052 (2011).

33. M. A. Dickson et al., Human keratinocytes that express hTERT and also bypass a p16INK4a-enforced mechanism that limits life span become immortal yet retain normal growth and differentiation characteristics. Molecular and cellular biology 20, 1436–1447 (2000).

34. L.-j. Zhang et al., Antimicrobial peptide LL37 and MAVS signaling drive interferon-β production by epidermal keratinocytes during skin injury. Immunity 45, 119–130 (2016).

35. E. Boelsma, M. C. Verhoeven, M. Ponec, Reconstruction of a human skin equivalent using a spontaneously transformed keratinocyte cell line (HaCaT). Journal of investigative dermatology 112, 489–498 (1999).

36. Y.-A. Cao et al., Gene expression profiling in pachyonychia congenita skin. Journal of dermatological science 77, 156–165 (2015).

37. O. Duverger, M. A. Cross, F. J. Smith, M. I. Morasso, Enamel anomalies in a pachyonychia congenita patient with a mutation in KRT16. Journal of Investigative Dermatology 139, 238–241 (2019).

38. F. J. Smith, et al., Pachyonychia congenita. (2017).

39. M. Wawersik, P. A. Coulombe, Forced expression of keratin 16 alters the adhesion, differentiation, and migration of mouse skin keratinocytes. Molecular biology of the cell 11, 3315–3327 (2000).

40. R. Vassar, P. A. Coulombe, L. Degenstein, K. Albers, E. Fuchs, Mutant keratin expression in transgenic mice causes marked abnormalities resembling a human genetic skin disease. Cell 64, 365–380 (1991).

41. A. J. Sadler, B. R. Williams, Interferon-inducible antiviral effectors. Nature reviews immunology 8, 559–568 (2008).

42. C.-P. Liao, E. Tchegnon, L. Q. Le, Double-stranded RNA sensing determines epithelial cell identity. Journal of Investigative Dermatology 139, 17–19 (2019).

43. M. N. H. Malik et al., Congenital deficiency reveals critical role of ISG15 in skin homeostasis. The Journal of Clinical Investigation 132, (2022).

44. P. F. Dos Santos, D. S. Mansur, Beyond ISGlylation: functions of free intracellular and extracellular ISG15. Journal of Interferon & Cytokine Research 37, 246–253 (2017).

45. L. Van Der Fits et al., Imiquimod-induced psoriasis-like skin inflammation in mice is mediated via the IL-23/IL-17 axis. The Journal of Immunology 182, 5836–5845 (2009).

46. A. Walter et al., Aldara activates TLR7-independent immune defence. Nature communications 4, 1560 (2013).

47. E. Sezer et al., Diagnostic utility of Ki-67 and Cyclin D1 immunostaining in differentiation of psoriasis vs. other psoriasiform dermatitis. Dermatology Practical & Conceptual 5, 7 (2015).

48. Y.-Z. Huang et al., OAS1, OAS2, and OAS3 contribute to epidermal keratinocyte proliferation by regulating cell cycle and augmenting IFN-1‒Induced jak1‒signal transducer and activator of transcription 1 phosphorylation in psoriasis. Journal of Investigative Dermatology 142, 2635–2645. e2639 (2022).

49. M. C. Dickson et al., A model of skin inflammation in humans leads to a rapid and reproducible increase in the interferon response signature: a potential translational model for drug development. Inflammation Research 64, 171–183 (2015).

50. W. R. Swindell et al., Imiquimod has strain-dependent effects in mice and does not uniquely model human psoriasis. Genome medicine 9, 1–21 (2017).

51. J. P. Smits et al., Immortalized N/TERT keratinocytes as an alternative cell source in 3D human epidermal models. Scientific reports 7, 11838 (2017).

52. M. K. Sarkar et al., Keratinocytes sense and eliminate CRISPR DNA through STING/IFN-κ activation and APOBEC3G induction. The Journal of clinical investigation 133, (2023).

53. R. B. Seth, L. Sun, C.-K. Ea, Z. J. Chen, Identification and characterization of MAVS, a mitochondrial antiviral signaling protein that activates NF-κB and IRF3. Cell 122, 669–682 (2005).

54. T. Kawai, S. Akira, Signaling to NF-κB by Toll-like receptors. Trends in molecular medicine 13, 460–469 (2007).

55. Y.-C. Perng, D. J. Lenschow, ISG15 in antiviral immunity and beyond. Nature Reviews Microbiology 16, 423–439 (2018).

56. S. D. Der, A. Zhou, B. R. Williams, R. H. Silverman, Identification of genes differentially regulated by interferon α, β, or γ using oligonucleotide arrays. Proceedings of the National Academy of Sciences 95, 15623–15628 (1998).

57. S. Ning, J. Pagano, G. Barber, IRF7: activation, regulation, modification and function. Genes & Immunity 12, 399–414 (2011).

58. R. Lin, C. Heylbroeck, P. M. Pitha, J. Hiscott, Virus-dependent phosphorylation of the IRF-3 transcription factor regulates nuclear translocation, transactivation potential, and proteasome-mediated degradation. Molecular and cellular biology, (1998).

59. F. Christian, E. L. Smith, R. J. Carmody, The regulation of NF-κB subunits by phosphorylation. Cells 5, 12 (2016).

60. J. Keller et al., LL37 complexed to double-stranded RNA induces RIG-I-like receptor signalling and Gasdermin E activation facilitating IL-36γ release from keratinocytes. Cell Death & Disease 16, 198 (2025).

61. A. K. Perry, E. K. Chow, J. B. Goodnough, W.-C. Yeh, G. Cheng, Differential requirement for TANK-binding kinase-1 in type I interferon responses to toll-like receptor activation and viral infection. The Journal of experimental medicine 199, 1651–1658 (2004).

62. H. M. Liu et al., The mitochondrial targeting chaperone 14-3-3ε regulates a RIG-I translocon that mediates membrane association and innate antiviral immunity. Cell host & microbe 11, 528–537 (2012).

63. F. Madeira et al., 14-3-3-Pred: improved methods to predict 14-3-3-binding phosphopeptides. Bioinformatics 31, 2276–2283 (2015).

64. D. K. Morrison, The 14-3-3 proteins: integrators of diverse signaling cues that impact cell fate and cancer development. Trends in cell biology 19, 16–23 (2009).

65. J. Liao, M. B. Omary, 14-3-3 proteins associate with phosphorylated simple epithelial keratins during cell cycle progression and act as a solubility cofactor. The Journal of cell biology 133, 345–357 (1996).

66. R. A. Mariani, S. Paranjpe, R. Dobrowolski, G. F. Weber, 14-3-3 targets keratin intermediate filaments to mechanically sensitive cell–cell contacts. Molecular biology of the cell 31, 930–943 (2020).

67. X. Pan, L. A. Kane, J. E. Van Eyk, P. A. Coulombe, Type I keratin 17 protein is phosphorylated on serine 44 by p90 ribosomal protein S6 kinase 1 (RSK1) in a growth-and stress-dependent fashion. Journal of Biological Chemistry 286, 42403–42413 (2011).

68. H. Zhu et al., RIG-I antiviral signaling drives interleukin-23 production and psoriasis-like skin disease. EMBO molecular medicine 9, 589–604 (2017).

69. F. Hou et al., MAVS forms functional prion-like aggregates to activate and propagate antiviral innate immune response. Cell 146, 448–461 (2011).

70. Y. Renert-Yuval et al., Biomarkers in atopic dermatitis—a review on behalf of the International Eczema Council. Journal of Allergy and Clinical Immunology 147, 1174–1190.e1171 (2021).

71. S. Gupta et al., 14-3-3 scaffold proteins mediate the inactivation of trim25 and inhibition of the type I interferon response by herpesvirus deconjugases. PLoS pathogens 15, e1008146 (2019).

72. S. Gupta, P. Ylä-Anttila, T. Sandalova, A. Achour, M. G. Masucci, Interaction with 14-3-3 correlates with inactivation of the RIG-I signalosome by herpesvirus ubiquitin deconjugases. Frontiers in Immunology 11, 437 (2020).

73. P. Awasthi, D. Kumar, S. Hasan, Role of 14-3-3 protein family in the pathobiology of EBV in immortalized B cells and Alzheimer’s disease. Frontiers in Molecular Biosciences 11, 1353828 (2024).

74. G. A. Hile, J. E. Gudjonsson, J. M. Kahlenberg, The influence of interferon on healthy and diseased skin. Cytokine 132, 154605 (2020).

75. N. H. Gopee et al., A prenatal skin atlas reveals immune regulation of human skin morphogenesis. Nature, 1–11 (2024).

76. S. Tilwani, K. Gandhi, S. N. Dalal, 14-3-3ε conditional knockout mice exhibit defects in the development of the epidermis. FEBS letters, (2024).

77. K. M. Bernot, P. A. Coulombe, K. M. McGowan, Keratin 16 expression defines a subset of epithelial cells during skin morphogenesis and the hair cycle. Journal of investigative dermatology 119, 1137–1149 (2002).

78. J. C. Lessard, P. A. Coulombe, Keratin 16–null mice develop palmoplantar keratoderma, a hallmark feature of pachyonychia congenita and related disorders. Journal of investigative dermatology 132, 1384–1391 (2012).

79. R. K. Thimmulappa et al., Nrf2 is a critical regulator of the innate immune response and survival during experimental sepsis. The Journal of clinical investigation 116, 984–995 (2016).

80. S. Han et al., NF-E2–related factor 2 regulates interferon receptor expression and alters macrophage polarization in lupus. Arthritis & Rheumatology 72, 1707–1720 (2020).

81. I. A. Buskiewicz et al., Reactive oxygen species induce virus-independent MAVS oligomerization in systemic lupus erythematosus. Science signaling 9, ra115–ra115 (2016).

82. R. Zhou et al., dsRNA sensing induces loss of cell identity. Journal of Investigative Dermatology 139, 91–99 (2019).

83. A. Batycka-Baran, J. Maj, R. Wolf, J. Szepietowski, The new insight into the role of antimicrobial proteins-alarmins in the immunopathogenesis of psoriasis. Journal of immunology research 2014, 628289 (2014).

84. M. Ma, W. Jiang, R. Zhou, DAMPs and DAMP-sensing receptors in inflammation and diseases. Immunity 57, 752–771 (2024).

85. K. Pennington, T. Chan, M. Torres, J. Andersen, The dynamic and stress-adaptive signaling hub of 14-3-3: emerging mechanisms of regulation and context-dependent protein–protein interactions. Oncogene 37, 5587–5604 (2018).

86. J. M. Kahlenberg, M. J. Kaplan, Little peptide, big effects: the role of LL-37 in inflammation and autoimmune disease. The Journal of Immunology 191, 4895–4901 (2013).

87. M. Wawersik, R. D. Paladini, E. Noensie, P. A. Coulombe, A proline residue in the α-helical rod domain of type I keratin 16 destabilizes keratin heterotetramers. Journal of Biological Chemistry 272, 32557–32565 (1997).

88. K. M. Bernot, C.-H. Lee, P. A. Coulombe, A small surface hydrophobic stripe in the coiled-coil domain of type I keratins mediates tetramer stability. The Journal of cell biology 168, 965–974 (2005).

89. C.-H. Lee, P. A. Coulombe, Self-organization of keratin intermediate filaments into cross-linked networks. Journal of Cell Biology 186, 409–421 (2009).

90. S. Sankar et al., A novel role for keratin 17 in coordinating oncogenic transformation and cellular adhesion in Ewing sarcoma. Molecular and cellular biology, (2013).

91. N.-O. Ku, S. Michie, E. Z. Resurreccion, R. L. Broome, M. B. Omary, Keratin binding to 14-3-3 proteins modulates keratin filaments and hepatocyte mitotic progression. Proceedings of the National Academy of Sciences 99, 4373–4378 (2002).

92. J. Basset, L. Marchal, A. Hovnanian, EGFR signaling is overactive in pachyonychia congenita: effective treatment with oral erlotinib. Journal of Investigative Dermatology 143, 294–304. e298 (2023).

93. D. P. McLornan, J. E. Pope, J. Gotlib, C. N. Harrison, Current and future status of JAK inhibitors. The lancet 398, 803–816 (2021).

94. K. Phan, D. Sebaratnam, JAK inhibitors for alopecia areata: a systematic review and meta-analysis. Journal of the European Academy of Dermatology and Venereology 33, 850–856 (2019).

95. D. J. Wallace et al., Baricitinib for systemic lupus erythematosus: a double-blind, randomised, placebo-controlled, phase 2 trial. The Lancet 392, 222–231 (2018).

96. K. Papp et al., Tofacitinib, an oral Janus kinase inhibitor, for the treatment of chronic plaque psoriasis: results from two randomized, placebo-controlled, phase III trials. British Journal of Dermatology 173, 949–961 (2015).

97. H. Bachelez et al., Tofacitinib versus etanercept or placebo in moderate-to-severe chronic plaque psoriasis: a phase 3 randomised non-inferiority trial. The Lancet 386, 552–561 (2015).

98. E. L. Simpson et al., Efficacy and safety of abrocitinib in adults and adolescents with moderate-to-severe atopic dermatitis (JADE MONO-1): a multicentre, double-blind, randomised, placebo-controlled, phase 3 trial. The Lancet 396, 255–266 (2020).

99. D. Rosmarin et al., Ruxolitinib cream for treatment of vitiligo: a randomised, controlled, phase 2 trial. The Lancet 396, 110–120 (2020).

100. E. A. O’Toole et al., Pachyonychia congenita: a research agenda leading to new therapeutic approaches. Journal of Investigative Dermatology 144, 748–754 (2024).

101. M. L. Kerns, J. M. Hakim, A. Zieman, R. G. Lu, P. A. Coulombe, Sexual dimorphism in response to an NRF2 inducer in a model for pachyonychia congenita. Journal of Investigative Dermatology 138, 1094–1100 (2018).

102. A. Zieman, P. A. Coulombe, The keratin 16 null phenotype is modestly impacted by genetic strain background in mice. Experimental dermatology 27, 672–674 (2018).

103. J. B. Cheng et al., Transcriptional programming of normal and inflamed human epidermis at single-cell resolution. Cell reports 25, 871–883 (2018).

104. F. Ma et al., Single cell and spatial sequencing define processes by which keratinocytes and fibroblasts amplify inflammatory responses in psoriasis. Nature communications 14, 3455 (2023).

105. J. E. Gudjonsson et al., Contribution of plasma cells and B cells to hidradenitis suppurativa pathogenesis. JCI insight 5, (2020).

106. H. Mi, A. Muruganujan, J. T. Casagrande, P. D. Thomas, Large-scale gene function analysis with the PANTHER classification system. Nature protocols 8, 1551–1566 (2013).

107. D. Croft et al., Reactome: a database of reactions, pathways and biological processes. Nucleic acids research 39, D691–D697 (2010).

108. M. Dai et al., Evolving gene/transcript definitions significantly alter the interpretation of GeneChip data. Nucleic acids research 33, e175–e175 (2005).

109. M. E. Ritchie et al., Empirical array quality weights in the analysis of microarray data. BMC bioinformatics 7, 1–16 (2006).

110. G. K. Smyth, Linear models and empirical bayes methods for assessing differential expression in microarray experiments. Statistical applications in genetics and molecular biology 3, (2004).

111. P. Boukamp et al., Normal keratinization in a spontaneously immortalized aneuploid human keratinocyte cell line. The Journal of cell biology 106, 761–771 (1988).

112. D. Mellacheruvu et al., The CRAPome: a contaminant repository for affinity purification– mass spectrometry data. Nature methods 10, 730–736 (2013).

113. H. Choi et al., SAINT: probabilistic scoring of affinity purification–mass spectrometry data. Nature methods 8, 70–73 (2011).

114. D. Szklarczyk et al., The STRING database in 2023: protein–protein association networks and functional enrichment analyses for any sequenced genome of interest. Nucleic acids research 51, D638–D646 (2023).

115. B. Xu et al., A critical role for IFN-β signaling for IFN-κ induction in keratinocytes. Frontiers in Lupus 2, 1359714 (2024).

116. C. A. Schneider, W. S. Rasband, K. W. Eliceiri, NIH Image to ImageJ: 25 years of image analysis. Nature methods 9, 671–675 (2012).

117. D. R. Stirling et al., CellProfiler 4: improvements in speed, utility and usability. BMC bioinformatics 22, 1–11 (2021).

