## Supplementary material for "Keratin 16 spatially inhibits type I interferon responses in stressed and diseased skin": Cohen et al. SUPPL MATERIALS all (Oct 2025)

### List of elements

**Supplemental Figure 1.** Spatial RNAseq and indirect immunofluorescence in human psoriatic skin.

**Supplemental Figure 2.** Additional data from IMQ-treated back skin in WT and Krt16 null mice.

**Supplemental Figure 3.** Additional data from N-TERT keratinocyte cultures, post-confluence.

**Supplemental Figure 4.** Inhibitor and rescue assays in keratinocyte cultures.

**Supplemental Table 1.** List of proteins co-immunoprecipitating with transfected, Flag-tagged K16 in HaCaT keratinocytes in ex vivo culture (SAINT>0.8, sorted according to SAINT score).

**Supplemental Table 2.** List of antibodies used in this study (Cohen et al.).

**Supplemental Table. 3.** information on animals treated with ruxolitinib.

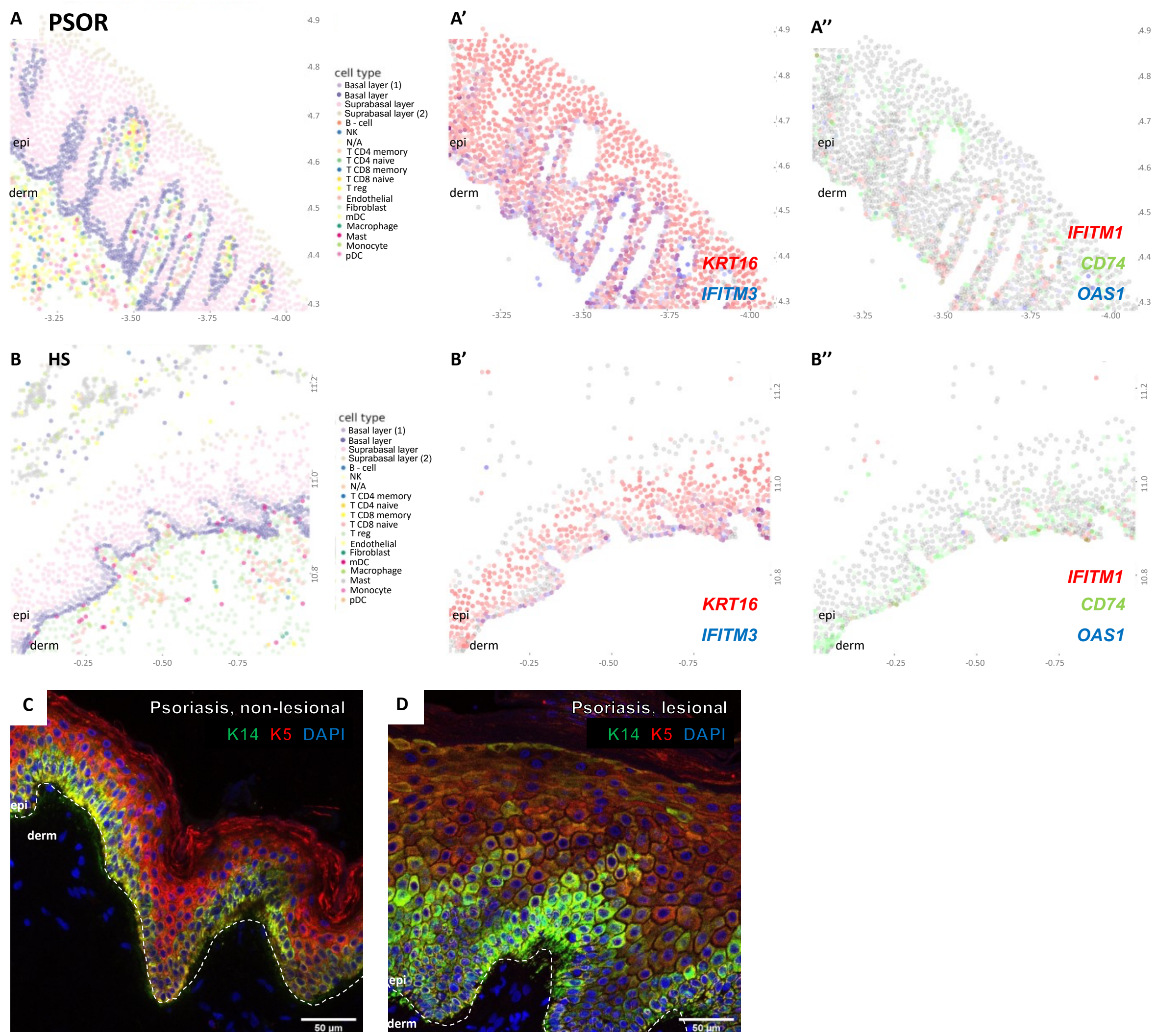


**Supplemental Figure 1. Spatial RNAseq and indirect immunofluorescence in human psoriatic skin.** (A-A'') Spatial RNAseq analysis of sections of lesional skin from individuals with psoriasis. Images depict the different cell types identified (A), expression of *KRT16* (red) and *IFITM3* (blue) in (A'), and of *IFITM1* (red), *CD74* (green) and *OAS1* (blue) in (A''). (B-B'') Spatial RNAseq analysis of lesional skin from individuals with hidradenitis suppurativa (HS). Images depict the different cell types identified (B), expression of *KRT16* (red) and IFITM3 (blue (B'), and of *IFITM1* (red), *CD74* (green) and *OAS1* (blue) (B''). (C-D) Indirect immunofluorescence for K14 (green), K5 (red) and staining for nuclei (DAPI, blue) in tissue sections non-lesional (C) or lesional (D) skin of individuals with psoriasis. Dashed white lines depict the dermo-epidermal interface.

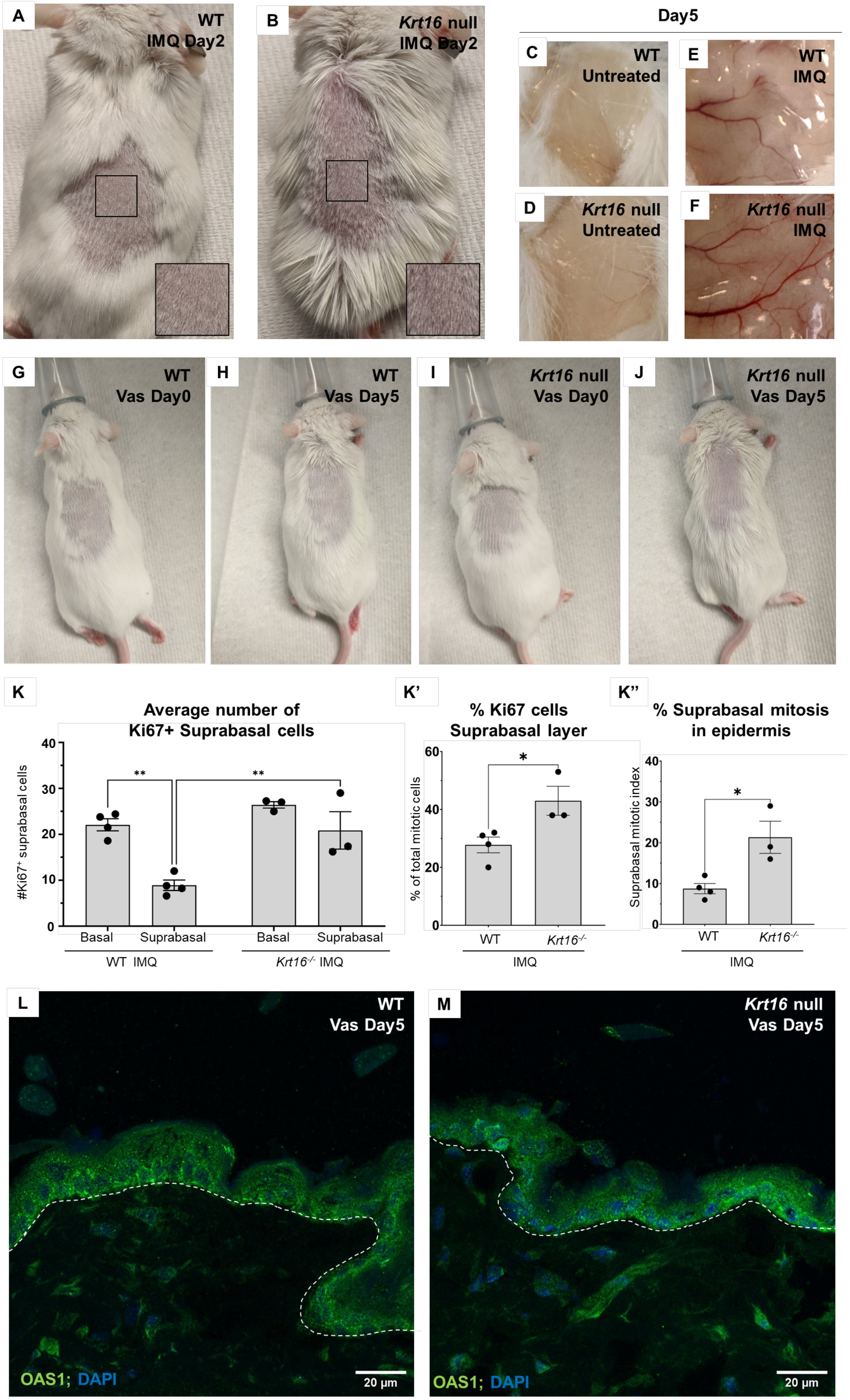

A
WT
IMQ Day2
B
Krt16 null
IMQ Day2
Day5
C
WT
Untreated
E
WT
IMQ
D
Krt16 null
Untreated
F
Krt16 null
IMQ
G
WT
Vas Day0
H
WT
Vas Day5
I
Krt16 null
Vas Day0
J
Krt16 null
Vas Day5
K
Average number of
Ki67+ Suprabasal cells
#Ki67+ suprabasal cells
40
30
20
10
0
**
**
Basal
Suprabasal
Basal
Suprabasal
WT IMQ
Krt16-/- IMQ
K'
% Ki67 cells
Suprabasal layer
% of total mitotic cells
60
40
20
0
*
WT
Krt16-/-
IMQ
K''
% Suprabasal mitosis
in epidermis
Suprabasal mitotic index
40
30
20
10
0
*
WT
Krt16-/-
IMQ
L
WT
Vas Day5
OAS1; DAPI
20 µm
M
Krt16 null
Vas Day5
OAS1; DAPI
20 µm

**Supplemental Figure 2. Additional data from IMQ-treated back skin in WT and *Krt16* null mice.** (A-B) Images of back skin after 2 days of IMQ application in WT (A) and *Krt16* null (B) mice. (C-F) Images of dissected back skin from 5 d unshaved, untreated back skin area (C-D) or IMQ-treated site (E-F) in WT (C,E) or *Krt16* null (D,F) mice. (G-J) Images of regions of interest prior to treatment, or after 5 d of Vaseline application to back skin in WT (G-H) and *Krt16* null mice (I-J). (K-K'') Analysis of Ki67 immunostaining in IMQ-treated WT and *Krt16* null mice, reporting on Ki67-positive cell number (K), percentage of Ki67-positive cells in suprabasal layers (K') and percentage of suprabasal mitoses in total epidermis (K''). (L) Indirect immunofluorescence of OAS1 staining in 5-day Vaseline-treated WT (L) and *Krt16* null (M) mice. OAS1 (green), DAPI (blue). Dashed white lines depict the dermo-epidermal interface. One-way ANOVA with Tukey's multiple comparisons test was used to compare conditions in (K), and unpaired t-test was used to compare conditions in (K') and (K'').

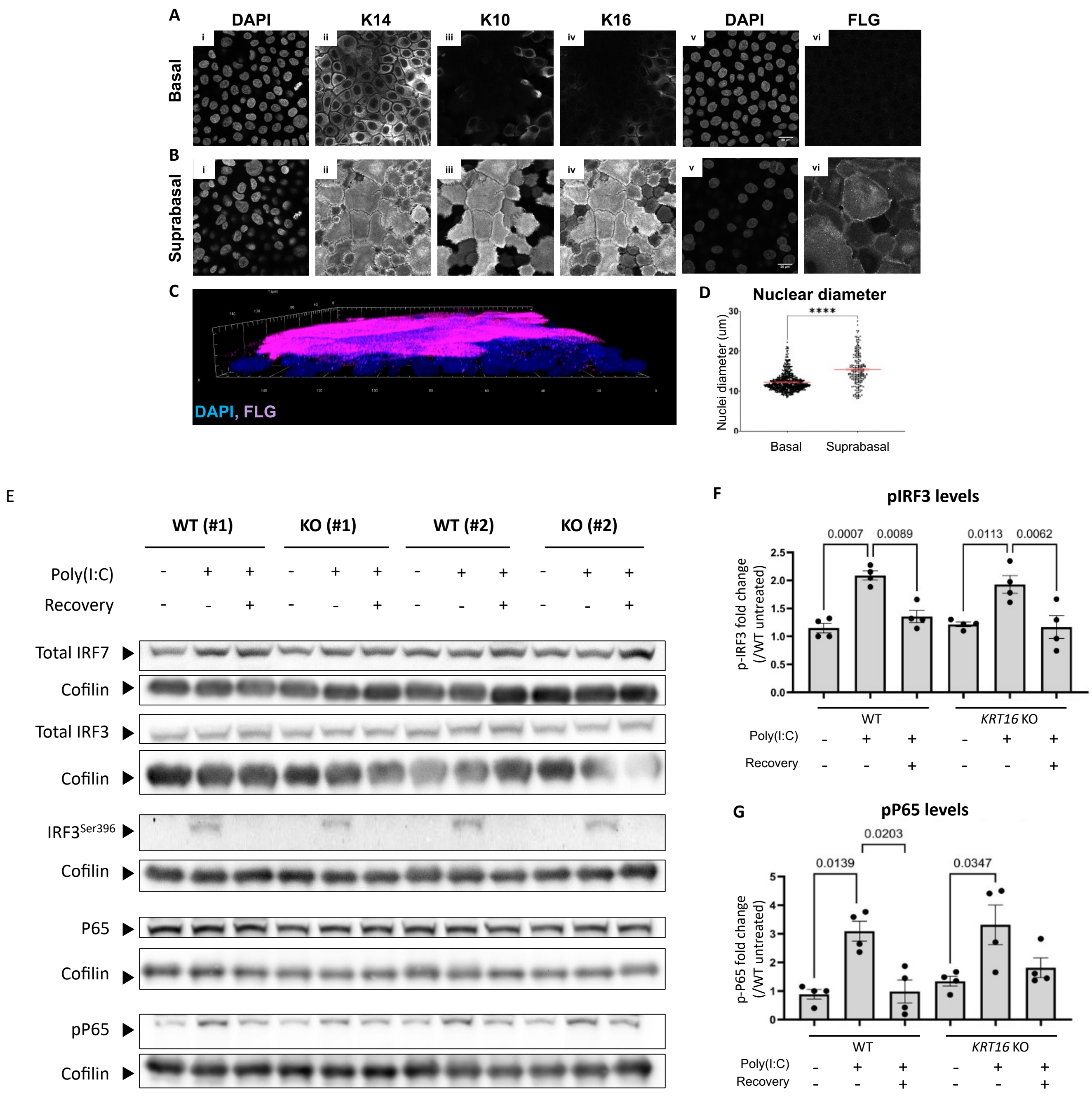


**Supplemental Figure 3. Additional data from N-TERT keratinocyte cultures, post-confluence.** (A-B) DAPI staining for nuclei (i and v), and indirect immunofluorescence for K14 (ii), K10 (iii), K16 (iv) and FLG (vi) in the basal layer (A) or suprabasal layer (B) of post-confluent N/TERT keratinocyte cultures. (C) 3D reconstruction of bilayered post-confluent N/TERT keratinocyte cultures stained for FLG (magenta) and DAPI (blue). (D) Quantification of nuclear diameter in the basal and suprabasal layers of N/TERT keratinocytes stained with DAPI. (E) Western blot analysis of total IRF3, phosphorylated IRF3, P65 and phosphorylated P65 in WT and *KRT16* KO post-confluent N/TERT cultures after (1) no treatment, (2) polyIC treatment without recovery and (3) polyIC treatment at 24h after recovery. Two biological replicates per condition are shown (WT#1, #2; KO#1, #2). Cofilin was used as a loading control. (F-G) Quantifications of p-IRF3 (F) and p-P65 (G) levels across polyIC treatment conditions in WT and *KRT16 KO* N/TERT cultures, post-confluence. Each dot represents a biological replicate. Unpaired t-test was used to compare conditions in (D). One-way ANOVA with Šídák's multiple comparisons test was used to compare conditions in (F-G).

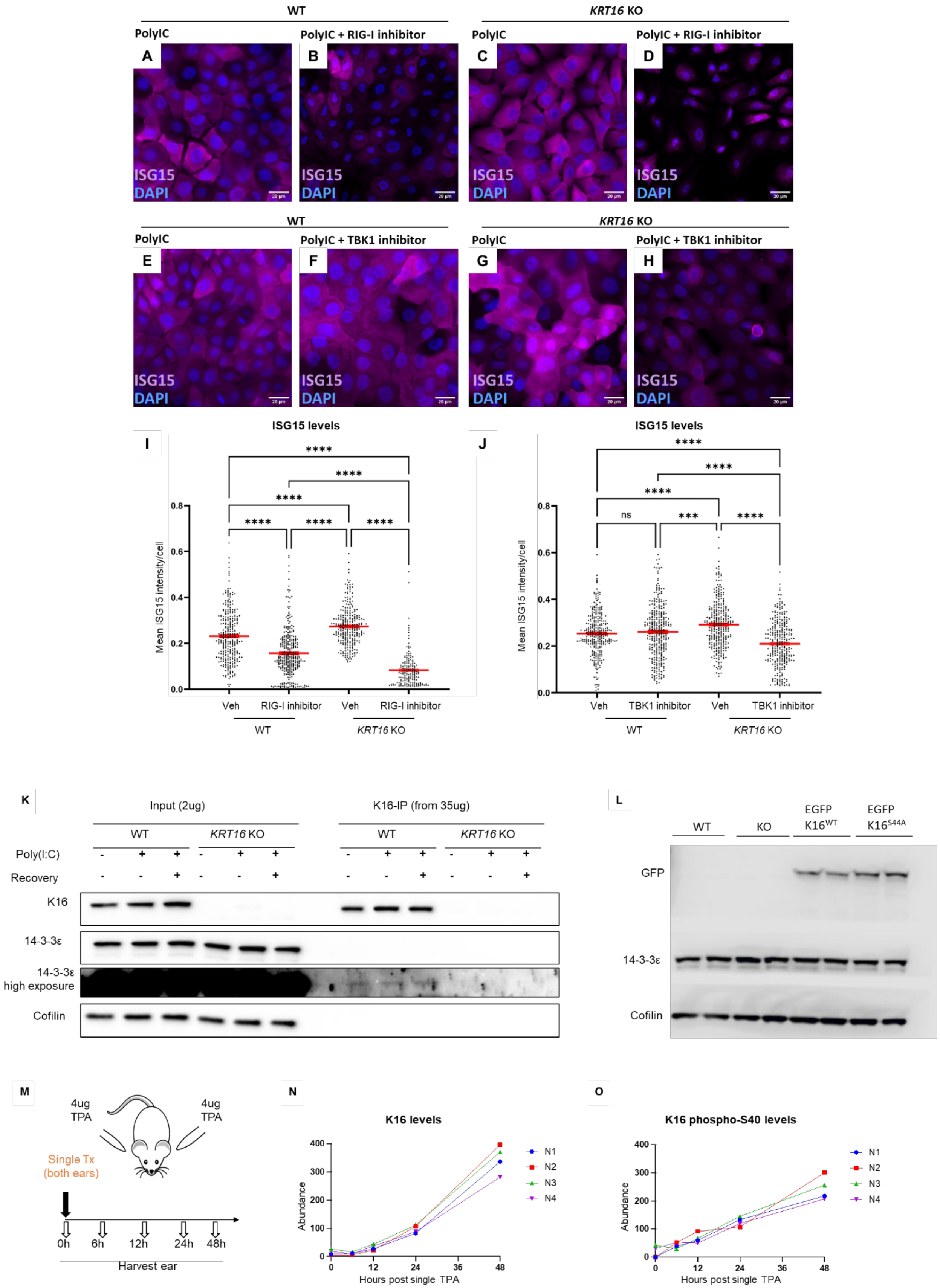

WT
KRT16 KO
PolyIC
PolyIC + RIG-I inhibitor
A
B
C
D
ISG15
DAPI
PolyIC + TBK1 inhibitor
E
F
G
H
I
ISG15 levels
Mean ISG15 intensity/cell
Veh
RIG-I inhibitor
J
TBK1 inhibitor
ns
K
Input (2ug)
K16-IP (from 35ug)
Poly(I:C)
Recovery
K16
14-3-3ε
14-3-3ε high exposure
Cofilin
L
KO
EGFP K16WT
EGFP K16S44A
GFP
M
4ug TPA
Single Tx (both ears)
0h
6h
12h
24h
48h
Harvest ear
N
K16 levels
Abundance
Hours post single TPA
N1
N2
N3
N4
O
K16 phospho-S40 levels

**Supplemental Figure 4. Inhibitor and rescue assays from keratinocyte cultures.** A-B) Max projections of DAPI staining for nuclei (blue) and indirect immunofluorescence for ISG15 (magenta) in the basal layer of poly(I:C) treated WT (A,B,E,F) or *KRT16 KO* (C,D,G,H) in absence or presence of RIG-I inhibitor (A-D) or TBK1 inhibitor (E-H). (I-J) Quantifications of inhibitor assays for RIG-I (I) or TBK1 (J). Points represent individual cells across 3 biological replicates. (K) Immunoprecipitation of endogenous K16 from WT or *KRT16 KO* cells. Input (2ug total) was used to validate genotype and assess for 14-3-3ε levels across conditions. K16 pulldown (IP from total 35 ug) was verified in the IP conditions, and longer exposure used to detect 14-3-3ε pulldown in the IP samples across conditions. (L) Whole cell lysates from two biological replicates of WT (lane 1-2), *KRT16 KO* (lane 3-4)and *KRT16 KO* expressing either EGFP-K16$^{WT}$ (lane 5-6) or EGFP-K16$^{S44A}$ (lane 7-8). 2 biological replicates are loaded per cell line and are blotted for GFP, 14-3-3ε and Cofilin loading control. (M) Illustration of approach used to generate a mass spectrometry phospho-proteomic profile of K16 at 0, 6, 12, 24 and 48 hours after a single TPA treatment of ear skin in WT mice (n= 3 animals). (N) Levels of K16 from MS analysis of TPA treated WT mouse ear. (O) Levels of phosphorylated S40 residue in K16 across time after TPA treatment.

**Supplemental Table 1. List of proteins co-immunoprecipating with transfected, Flag-tagged K16 in HaCaT keratinocytes in *ex vivo* culture (SAINT>0.8, sorted according to SAINT score)**

| Bait | Prey | GeneID | Quant | Control | Avg. SPC | FCA | FCB | SAINT |
|---|---|---|---|---|---|---|---|---|
| KRT16 | P08779 | KRT16 | 45\|46\|43 | 0\|0\|1 | 44.67 | 64.69 | 47.58 | 1 |
| KRT16 | P09913 | IFIT2 | 5\|22\|36 | 0\|3\|3 | 21.00 | 8.32 | 8.31 | 1 |
| KRT16 | P09914 | IFIT1 | 17\|28\|46 | 2\|5\|1 | 30.33 | 8.46 | 7.82 | 1 |
| KRT16 | O14879 | IFIT3 | 10\|32\|57 | 0\|9\|5 | 33.00 | 7.27 | 7.26 | 1 |
| KRT16 | P20591 | MX1 | 7\|35\|77 | 0\|0\|0 | 39.67 | 35.64 | 34.79 | 1 |
| KRT16 | Q9Y6K5 | OAS3 | 1\|14\|22 | 0\|3\|0 | 12.33 | 5.83 | 5.18 | 0.995 |
| KRT16 | Q96AZ6 | ISG20 | 8\|6\|11 | 0\|0\|0 | 8.33 | 14.77 | 11.48 | 0.985 |
| KRT16 | Q7Z2W4 | ZC3HAV1 | 0\|32\|34 | 0\|13\|6 | 22.00 | 2.92 | 1.42 | 0.975 |
| KRT16 | O14933 | UBE2L6 | 0\|10\|15 | 0\|1\|3 | 8.33 | 3.66 | 2.41 | 0.96 |
| KRT16 | O95786 | DDX58 | 1\|16\|20 | 0\|0\|0 | 12.33 | 11.24 | 9.87 | 0.955 |
| KRT16 | Q8IY21 | DDX60 | 0\|6\|21 | 0\|1\|0 | 9.00 | 5.39 | 3.46 | 0.945 |
| KRT16 | Q9BYX4 | IFIH1 | 0\|15\|20 | 0\|0\|0 | 11.67 | 9.71 | 5.80 | 0.925 |
| KRT16 | Q15646 | OASL | 0\|7\|19 | 0\|0\|0 | 8.67 | 6.92 | 4.54 | 0.92 |
| KRT16 | P29728 | OAS2 | 1\|8\|11 | 0\|0\|0 | 6.67 | 6.92 | 6.69 | 0.91 |
| KRT16 | P00973 | OAS1 | 2\|6\|15 | 0\|0\|0 | 7.67 | 8.26 | 8.17 | 0.895 |
| KRT16 | Q460N5 | PARP14 | 0\|5\|10 | 0\|1\|0 | 5.00 | 3.51 | 2.63 | 0.875 |
| KRT16 | Q9BQE5 | APOL2 | 0\|6\|7 | 0\|0\|1 | 4.33 | 3.52 | 2.67 | 0.855 |
| KRT16 | Q9Y3Z3 | SAMHD1 | 2\|6\|10 | 0\|2\|1 | 6.00 | 4.04 | 4.02 | 0.82 |

**Supplemental Table 1 (continued). List of proteins co-immunoprecipating with endogenous K16 in N/TERT keratinocytes**

| Gene Symbol | Accession | Abundance Untreated | Abundance PolyIC + 24h | Abundance Count Untreated | Abundance Count PolyIC + 24h | # PSMs | # Unique Peptides |
|---|---|---|---|---|---|---|---|
| KRT5 | P13647 | 11032635913 | 10728657149 | 76 | 69 | 850 | 36 |
| KRT14 | P02533 | 4762651893 | 4847602534 | 48 | 42 | 822 | 17 |
| KRT6A | P02538 | 4874954708 | 4930642575 | 40 | 36 | 705 | 4 |
| KRT6B | P04259 | 258974639.3 | 175707599 | 5 | 4 | 613 | 4 |
| DSP | P15924 | 1097079742 | 1094440648 | 160 | 131 | 591 | 148 |
| KRT17 | Q04695 | 6464756965 | 6540763594 | 63 | 58 | 589 | 27 |
| KRT16 | P08779 | 726834130.5 | 633679244.2 | 28 | 25 | 495 | 23 |
| KRT1 | P04264 | 2152528216 | 836119936 | 52 | 44 | 456 | 37 |
| KRT9 | P35527 | 1237232329 | 431567670.8 | 53 | 41 | 359 | 36 |
| PLEC | Q15149 | 393356355.9 | 421978653.6 | 133 | 106 | 345 | 141 |
| KRT10 | P13645 | 961276840.3 | 583801883 | 40 | 33 | 303 | 28 |
| KRT2 | P35908 | 333023154.4 | 103475854.5 | 31 | 20 | 275 | 24 |
| KRT15 | P19012 | 199412663.6 | 179890113.9 | 24 | 19 | 272 | 15 |
| KRT8 | P05787 | 492952469.2 | 533347896.9 | 34 | 29 | 227 | 27 |
| ALB | P02768 | 1115164203 | 1412350143 | 12 | 11 | 168 | 12 |
| KRT13 | P13646 | 93188593.5 | 59934720 | 2 | 1 | 166 | 2 |
| SFN | P31947 | 260566108.5 | 252524932.4 | 16 | 16 | 145 | 13 |
| KRT19 | P08727 | 427359988.3 | 382750129.8 | 2 | 2 | 139 | 1 |
| YWHAQ | P27348 | 753599636.3 | 714233740.2 | 23 | 22 | 133 | 13 |
| MYH9 | P35579 | 188002323.3 | 74635575.19 | 55 | 28 | 120 | 53 |
| YWHAZ | P63104 | 188459664 | 147034368.8 | 12 | 11 | 101 | 12 |
| KRT76 | Q01546 | 6458491.688 | 2054835.625 | 2 | 1 | 97 | 2 |
| KRT79 | Q5XKE5 | 28715014 | 64131056 | 1 | 1 | 86 | 1 |
| RPL18A | Q02543 | 198100543.3 | 207915014.8 | 12 | 10 | 77 | 12 |
| CRYBG1 | Q9Y4K1 | 101754345.1 | 70257099 | 29 | 20 | 75 | 32 |
| VIM | P08670 | 74450732.25 | 53869058.25 | 23 | 16 | 73 | 24 |
| ACTB | P60709 | 138348039.3 | 89336881.5 | 20 | 15 | 72 | 1 |
| YWHAE | P62258 | 63945440.81 | 57766353.5 | 8 | 9 | 72 | 9 |
| ACTG1 | P63261 | 591756.5313 | 913313.3125 | 2 | 2 | 72 | 1 |
| YWHAB | P31946 | 41957990.88 | 29030901.5 | 5 | 4 | 70 | 5 |
| YWHAG | P61981 | 38267393.13 | 23596960.69 | 9 | 5 | 66 | 7 |
| A2M | P01023 | 82409304.88 | 85184774.25 | 9 | 9 | 62 | 9 |
| RPL4 | P36578 | 78659924.75 | 67057407.75 | 17 | 12 | 59 | 16 |
| EEF1A1 | P68104 | 72430136.38 | 75138944.75 | 12 | 9 | 58 | 12 |
| EPPK1 | P58107 | 26150630.47 | 8779960.443 | 12 | 4 | 57 | 13 |
| ACTG2 | P63267 | 17059067.25 | 9364873 | 3 | 3 | 50 | 3 |
| YWHAH | Q04917 | 9113050.438 | 7140637 | 3 | 2 | 49 | 3 |
| TUBA1B | P68363 | 37940424.88 | 31475327.06 | 13 | 9 | 48 | 3 |
| GAPDH | P04406 | 40609290.88 | 36445295.69 | 13 | 10 | 48 | 14 |
| RPS2 | P15880 | 72544530.63 | 71632170 | 11 | 10 | 46 | 11 |
| PKP1 | Q13835 | 45158100.19 | 39933613.56 | 18 | 13 | 45 | 19 |
| TUBB | P07437 | 5589073.875 | 3608855 | 2 | 1 | 45 | 2 |
| KRT18 | P05783 | 30239254.81 | 25979950.75 | 8 | 6 | 44 | 10 |
| KRT77 | Q7Z794 | 149692238.9 | 59492108 | 1 | 1 | 44 | 1 |
| RPS3 | P23396 | 61416715.38 | 54557362.5 | 14 | 12 | 43 | 14 |
| RPS18 | P62269 | 61976755.88 | 54997218.38 | 9 | 9 | 43 | 8 |
| TUBB4B | P68371 | 51164832.13 | 39700398.25 | 15 | 10 | 42 | 1 |
| RPS4X | P62701 | 44066451.22 | 38823859.38 | 14 | 11 | 39 | 13 |
| RPL6 | Q02878 | 47080323.13 | 49472537.75 | 10 | 9 | 38 | 10 |
| TUBA4A | P68366 | 3944643.063 | 2100574.5 | 3 | 1 | 37 | 3 |
| FAM83H | Q6ZRV2 | 43473271.81 | 25483189.63 | 19 | 12 | 37 | 20 |
| PKP3 | Q9Y446 | 48803669.19 | 33601789.5 | 19 | 10 | 36 | 21 |
| RPS8 | P62241 | 36323521.13 | 36434812 | 7 | 6 | 33 | 7 |
| RPS16 | P62249 | 40889446.63 | 39881126.19 | 9 | 6 | 32 | 8 |

### Supplemental Table 2. List of antibodies used in this study (Cohen et al.).

| Antigen | Host species | Company and catalog number | Dilution |
|---|---|---|---|
| **For indirect immunofluorescence:** | | | |
| K14 | chicken | BioLegend #906004 | 1:1000 |
| K5 | rabbit | Biolegend #10956 | 1:500 |
| K16 #1275 (human sections) | rabbit | | 1:400 |
| K16 (N/TERT) | mouse | Santa Cruz #sc-377224 | 1:100 |
| Ly6g (Phycoerythrin conjugated) | rat | BioLegend #127607 | 1:100 |
| OAS1 | rabbit | Proteintech #14955-1-AP | 1:500 |
| Ki67 | rabbit | Cell Signaling #12202 | 1:400 |
| ISG15 | rabbit | Proteintech #15981-1-AP | 1:400 |
| pIRF7 | rabbit | Cell Signaling #12390S | 1:200 |
| RIG-I | rabbit | EMD Millipore #MABF297 | 1:400 |
| 14-3-3 epsilon | rabbit | Proteintech #11648-2-AP | 1:400 |
| MAVS (rodent) | rabbit | Cell signaling #4983S | 1:500 |
| chicken IgY (Alexa 647 conjugated) | goat | Abcam #150171 | 1:1000 |
| rabbit IgG (Alexa 647 conjugated) | goat | Jackson ImmunoResearch #111-607-003 | 1:400 |
| ms IgG (Alexa 647 conjugated) | goat | Jackson ImmunoResearch #115-605-146 | 1:400 |
| rabbit IgG (Alexa 594 conjugated) | goat | Thermo Fisher Scientific #A11037 | 1:1000 |
| ms IgG (Alexa 594 conjugated) | goat | Fisher Scientific #A11032 | 1:1000 |
| rabbit IgG (Alexa 488 conjugated) | goat | Abcam #150077 | 1:1000 |
| mouse IgG (Alexa 488 conjugated) | goat | Jackson ImmunoResearch #115-545-003 | 1:400 |
| **Proximity ligation assays (PLA)** | | | |
| K16 (N/TERT) | mouse | Santa Cruz #sc-377224 | 1:200 |
| 14-3-3 epsilon | rabbit | Proteintech #11648-2-AP | 1:500 |
| 14-3-3 pan | mouse | Santa Cruz #sc-1657 | 1:200 |
| RIG-I | rabbit | EMD Millipore #MABF297 | 1:500 |
| **For western blotting and IP:** | | | |
| K16 #1275 (human sections) | rabbit | | 1:1000 |
| K16 LL025 | mouse | Santa cruz #sc-53255 | 1:500 |
| Total IRF7 | rabbit | Proteintech #22392-1-AP | 1:1000 |
| p-IRF7 | rabbit | Cell Signaling #12390S | 1:1000 |
| Total IRF3 | rabbit | Cell Signaling #4302S | 1:1000 |
| p-IRF3 | rabbit | Cell Signaling #4947S | 1:1000 |
| Total P65 | rabbit | Thermo Scientific #14-6731-81 | 1:1000 |
| pP65 | rabbit | Cell Signaling #3033 | 1:1000 |
| ISG15 | rabbit | Proteintech #15981-1-AP | 1:1000 |
| Cofilin | rabbit | Cell Signaling #5175S | 1:1000 |
| HSP90 | rabbit | Cell Signaling #4874S | 1:1000 |
| rabbit IgG (HRP linked) | goat | Cell Signaling Technology #7074P2 | 1:3000 |
| mouse IgG (HRP linked) | goat | Cell Signaling Technology #7076P2 | 1:3000 |

**Supplemental Table. 3: information on animals treated with ruxolitinib**

| Mouse (ID) | Background | Genotype | Age | Sex | Condition | PPK Epidermis Thickness (avg um) |
|---|---|---|---|---|---|---|
| 3159 | C57/BL/6J | *Krt16* -/- | 12 weeks | F | Veh | 417.4 |
| | | | | | Ruxo | 363.8 |
| 3160 | C57/BL/6J | *Krt16* -/- | 12 weeks | M | Veh | 593.1 |
| | | | | | Ruxo | 374.2 |
| 3320 | C57/BL/6J | *Krt16* -/- | 14 weeks | M | Veh | 393.7 |
| | | | | | Ruxo | 243.9 |
| 3321 | C57/BL/6J | *Krt16* -/- | 14 weeks | M | Veh | 519.6 |
| | | | | | Ruxo | 536.7 |
| 3322 | C57/BL/6J | *Krt16* -/- | 14 weeks | M | Veh | 449.7 |
| | | | | | Ruxo | 285.2 |
| 3324 | C57/BL/6J | *Krt16* -/- | 14 weeks | F | Veh | 353.6 |
| | | | | | Ruxo | 254.0 |
| 3325 | C57/BL/6J | *Krt16* -/- | 14 weeks | F | Veh | 353.4 |
| | | | | | Ruxo | 319.5 |
| 3323 | C57/BL/6J | *Krt16* -/- | 14 weeks | M | Veh | 289.5 |
| | | | | | Ruxo | 192.7 |
| 3326 | C57/BL/6J | *Krt16* -/- | 14 weeks | F | Veh | 331.8 |
| | | | | | Ruxo | 194.8 |
| 3158 | C57/BL/6J | *Krt16* +/+ | 12 weeks | F | Veh | 87.0 |
| | | | | | Ruxo | 54.4 |
| 3161 | C57/BL/6J | *Krt16* +/+ | 12 weeks | M | Veh | 61.7 |
| | | | | | Ruxo | 61.5 |
| 3681 | C57/BL/6J | *Krt16* +/+ | 14 weeks | M | Veh | 78.3 |
| | | | | | Ruxo | 71.5 |
| 3682 | C57/BL/6J | *Krt16* +/+ | 14 weeks | M | Veh | 83.9 |
| | | | | | Ruxo | 81.5 |
| 3683 | C57/BL/6J | *Krt16* +/+ | 14 weeks | M | Veh | 70.9 |
| | | | | | Ruxo | 69.8 |
